# Paired and solitary ionocytes in the zebrafish olfactory epithelium

**DOI:** 10.1101/2024.11.08.620918

**Authors:** Julia Peloggia, King Yee Cheung, Mariela D. Petkova, Richard Schalek, Jonathan Boulanger-Weill, Yuelong Wu, Shuohong Wang, Nicholas J. van Hateren, Michał Januszewski, Viren Jain, Jeff W. Lichtman, Florian Engert, Tatjana Piotrowski, Tanya T. Whitfield, Suresh Jesuthasan

## Abstract

The sense of smell is generated by electrical currents that are influenced by the concentration of ions in olfactory sensory neurons and mucus. In contrast to the extensive morphological and molecular characterization of sensory neurons, there has been little description of the cells that control ion concentrations in the zebrafish olfactory system. Here, we report the molecular and ultrastructural characterization of zebrafish olfactory ionocytes. Transcriptome analysis suggests that the zebrafish olfactory epithelium contains at least three different ionocyte types, which resemble Na^+^/K^+^-ATPase-rich (NaR), H^+^-ATPase-rich (HR), and Na^+^/Cl^-^ cotransporter (NCC) cells, responsible for calcium, pH, and chloride regulation, respectively, in the zebrafish skin. In the olfactory epithelium, NaR-like and HR-like ionocytes are usually adjacent to one another, whereas NCC-like cells are usually solitary. The distinct subtypes are differentially distributed: NaR-like/HR-like cell pairs are found broadly within the olfactory epithelium, whereas NCC-like cells reside within the peripheral non-sensory multiciliated cell zone. Comparison of gene expression and serial-section electron microscopy analysis indicates that the NaR-like cells wrap around the HR-like cells and are connected to them by shallow tight junctions. The development of olfactory ionocyte subtypes is also differentially regulated, as pharmacological Notch inhibition leads to a loss of NaR-like and HR-like cells, but does not affect NCC-like ionocyte number. These results provide a molecular and anatomical characterization of olfactory ionocytes in a stenohaline freshwater teleost. The paired ionocytes suggest that both transcellular and paracellular transport regulate ion concentrations in the olfactory epithelium, while the solitary ionocytes may enable independent regulation of ciliary beating.

## Introduction

Olfaction is mediated by a combination of ion currents — an influx of calcium followed by an efflux of chloride — that is triggered by the binding of an odorant to its receptor on olfactory sensory neurons (OSNs) (Kleene, 1993; Reuter et al., 1998). The high concentration of chloride in the dendritic knobs and cilia of mammalian OSNs is achieved by uptake from the mucus by the NKCC1 (Slc12a2) co-transporter, which requires external sodium (Kaneko et al., 2004). Despite the relative robustness of transduction that is offered by the chloride current (Reisert and Reingruber, 2019), olfactory sensitivity is influenced by external ion concentrations (Selvaraj et al., 2012). The zebrafish, a freshwater fish that can be found in the wild in water with a moderate range of salinity (Spence et al., 2006), has provided fundamental insights into olfactory processing (Friedrich et al., 2004; Niessing and Friedrich, 2010). The morphological and molecular features of OSNs of this animal — for example the location and transduction machinery in ciliated and microvillous neurons — are well described (Hansen and Zeiske, 1993, 1998; Saraiva et al., 2015). However, it is unclear how ion composition in the zebrafish olfactory epithelium and mucus is regulated.

Ionocytes are mitochondria-rich cells that can transport ions intracellularly against their concentration gradient. In freshwater teleosts, ionocytes in the skin and gills actively absorb ions from the external environment, compensating for passive water gain (reviewed in (Hwang and Chou, 2013; Guh et al., 2015)). In marine teleosts, ionocytes in the kidneys and gills actively secrete excess ions absorbed from seawater. Recently, with the advent of single-cell sequencing technologies, ionocytes have been identified based on gene expression in several tissues including the olfactory epithelium in humans and mice (Casellas et al., 2023; Ualiyeva et al., 2024), inner ear of mice (Honda et al., 2017) and the lateral line neuromasts in zebrafish (Peloggia et al., 2021). In the neuromasts, as in the gill and skin of fish, a subset of ionocytes were found in a complex; the presence of shallow tight junctions in such complexes (Hwang, 1988b) is thought to provide an alternative paracellular pathway for ion movement, for example by coupling Na^+^ secretion to Cl^-^ efflux via the CFTR (cystic fibrosis transmembrane conductance regulator) channel in the gill of *Fundulus* in seawater (Cozzi et al., 2015) .

The existence of ionocytes in the olfactory epithelium of fish was first suggested in 1972 based on light and electron microscopy of Baltic Sea trout (Bertmar, 1972). The cells, termed “labyrinth cells”, were characterized by an abundance of mitochondria as well as smooth endoplasmic reticulum, and were proposed to be functionally equivalent to “chloride cells” in the gill. A transmission electron microscopy (TEM) and scanning electron microscopy (SEM) study in 2001 confirmed the presence of cells with a similar appearance in the olfactory epithelium of seven freshwater fish species, and noted their distinct morphology: an apical surface with microvilli-like projections and occasional invaginations of the cell membrane (Ruzhinskaya et al., 2001). Labyrinth cells were also recently identified in the freshwater goby, where they were described as having a globular appearance in scanning electron micrographs (Ghosh, 2020). However, little is known about the molecular characteristics of these putative ionocytes, and it is not known if they exist in isolation or as complexes.

In mammals and frogs, the winged helix/forkhead transcription factor Foxi1 is required for ionocyte specification, and thus provides a useful marker for ionocytes. In zebrafish, the Foxi1 orthologs *foxi3a* and *foxi3b* regulate ionocyte development, and loss of *foxi3a* leads to complete loss of ionocytes (Hsiao et al., 2007; Jänicke et al., 2007). Based on their *foxi3a* expression, several classes of epidermal and gill ionocytes in the zebrafish were discovered, including H^+^-ATPase-rich (HR) ionocytes, which secrete protons, take up sodium, and excrete ammonium ions; Na^+^/-K^+^-ATPase-rich (NaR) ionocytes, which take up calcium ions; Na^+^/Cl^-^ co-transporter-expressing (NCC) ionocytes, which take up sodium and chloride ions; and K^+^-secreting (KS) ionocytes, which secrete potassium ions (reviewed in (Hwang, 2009; Hwang and Chou, 2013; Guh et al., 2015)). The different classes of ionocytes possess distinct gene expression profiles (Guh et al., 2015; Peloggia et al., 2021); distinguishing markers include *trpv6* (for NaR (Pan et al., 2005)), *ceacam1* (for HR (Kowalewski et al., 2021)), and *slc12a10.2* (for NCC (Wang et al., 2009; Shih et al., 2023)). Here, we use transcriptomic data, *in situ* hybridization and serial-section electron microscopy (ssEM) to characterize the different subtypes of ionocytes in the zebrafish olfactory epithelium.

## Materials and Methods

### Zebrafish husbandry

Zebrafish strains used in this study were AB and ABTL strain wild types, *nacre* (*mitfa^-/-^*) (Lister et al., 1999), *oval* (*ift88^tz288b^*) (Tsujikawa and Malicki, 2004), *Tg(tp1bglobin:EGFP)^um14^* (Parsons et al., 2009), *Tg(krtt1c19e:lyn-tdTomato)^sq16^*(Lee et al., 2014), *Tg(dld:hist2h2l-EGFP)^psi84^* (Kraus et al., 2022), and *Tg(-8.0cldnb:lyn-EGFP)^zf106Tg^* (Haas and Gilmour, 2006). Adult zebrafish were kept on a 10-hour dark/14-hour light cycle at 28.5°C and spawned by pair-mating or marbling. Eggs were collected and staged according to standard protocols, and raised either in 0.5× embryo E2 medium (7.5 mM NaCl, 0.25 mM KCl, 0.5 mM MgSO_4_, 75 mM KH_2_PO_4_, 25 mM Na_2_HPO_4_, 0.5 mM CaCl_2_, 0.5 mg/L NaHCO_3_, pH = 7.4) or 1× E3 medium (5 mM NaCl, 0.17 mM KCl, 0.33 mM CaCl_2_, 0.33 mM MgSO_4_, with 0.0001% methylene blue at early stages) at 28.5°C. Larvae were anesthetized with 0.5 mM tricaine methanesulfonate (MS222) at pH 7.

### Dissection of adult olfactory organs

Adult ABTL wild-type strain zebrafish were culled on ice and fixed in 4% paraformaldehyde (PFA) in 1× phosphate-buffered saline (PBS) overnight at 4°C. Adult zebrafish were transferred to a SYLGARD 184-coated (Dow Corning) Petri dish containing PBS, and olfactory organs were dissected out using Dumont #5SF forceps (Fine Science Tools). Olfactory organs were washed in PBS before proceeding with staining protocols.

### Single-cell RNA sequencing (scRNA-seq) analysis

Analysis was carried out on published datasets generated from dissected adult zebrafish olfactory organs (Kraus et al., 2022). All four datasets from PBS-treated (control) olfactory epithelia were downloaded with sratoolkit (version 3.1.1). Reads were de-multiplexed and aligned to version Ensembl GRCz11 (danRer11.Ens_106) of the zebrafish genome using the CellRanger (version 7.1.0) pipeline. Subsequent analysis of the UMI count matrix was performed using Seurat (version 5.1.0) (Satija et al., 2015; Butler et al., 2018; Stuart et al., 2019; Hao et al., 2021, 2024) in R version 4.3.3. Initial quality control filtered out genes expressed in fewer than three cells and cells with fewer than 200 genes. Further quality control was performed to exclude cells with more than 20000 UMIs or more than 10% mitochondrial content. The four resulting Seurat objects were combined with the function *merge()*, and cluster markers were identified with the function *FindAllMarkers()*. Dimensional reduction was performed (UMAP), and final clusters were obtained with 30 dimensions and resolution of 0.5.

Daniocell data were downloaded from the taste/olfactory subset (Sur et al., 2023); no additional processing or clustering was performed. Ionocytes were identified based on their expression of *foxi3b*. For solitary ionocytes, a dataset from larvae was used. Olfactory epithelial cells were subset based on cluster expressions of *ompb* and *trpc2a/2b*. Solitary olfactory ionocytes were then identified and manually clustered based on their *foxi3b* expression, using the function subset (object, subset = *foxi3b* > 0). Markers were then subsequently identified with the function *FindAllMarkers()*. Feature plots for all datasets were made with the function *FeaturePlot_scCustom()* from the package scCustomize.

### Hybridization chain reaction RNA-fluorescence *in situ* hybridization (HCR RNA-FISH)

HCR RNA-FISH was performed on 5 days post-fertilization (dpf) stage (unless stated otherwise) *nacre* wild-type or *Tg(tp1bglobin:EGFP)* transgenic larvae following “HCR RNA-FISH protocol for whole-mount zebrafish embryos and larvae (*Danio rerio*)” provided by Molecular Instruments, or adapted with acetone-based permeabilization (Peloggia et al., 2021). The probe sets used in this project were *ceacam1*-B1 (accession #: NM_001113794), *trpv6*-B1 and B2 (accession #: NM_001001849), *foxi3a*-B2 (accession #: NM_198917.2), *foxi3b*-B3 and B4 (accession #: NM_198918), *slc12a10.2*-B1 (accession #: NM_001045001.1), *slc4a1b*-B1 (accession #: NM_001168266.1), *slc9a3.2*-B4 (accession #: NM_001113479.1), *chrd*-B2 (accession #: NM_130973.3), and *hepacam2-*B2 (accession #: NM_001245085.1). The amplifiers used were B1-488, B1-647, B2-546, B2-594, B3-546, B3-647, and B4-488 (Molecular Instruments). To label all nuclei, larvae were incubated in 5 µg/mL DAPI for 30 minutes at room temperature. All samples were stored in PBS at 4°C before imaging. The above standard HCR RNA-FISH protocol for larvae was modified for staining on dissected olfactory organs of adult ABTL fish. The proteinase K treatment step was adjusted to incubation in 30 µg/ml of proteinase K for 30 minutes. The remainder of the protocol remained the same.

### Confocal imaging

Fixed zebrafish larvae and olfactory organs were mounted in 1–2% low melting point (LMP) agarose in PBS in glass-bottomed dishes, with larvae mounted in a dorsal view. Samples were imaged on either a Zeiss LSM 800 attached to an upright microscope with a W Plan Apochromat 40×/1.0 DIC M27 water-dipping objective, a Zeiss LSM 880 Airyscan confocal microscope equipped with a Plan-Apochromat 20×/0.8 M27 air objective, a Zeiss LSM 980 Airyscan2 confocal microscope equipped with a Plan-Apochromat 10×/0.45 air objective and an LCI Plan-Apochromat 40×/1.2 water objective acquired in Airyscan SR-4Y mode, or a Nikon Ti2 Yokogawa CSU-W1 spinning disk head equipped with a Hamamatsu Orca Fusion sCMOS. Objective lenses used on the Nikon microscope were CFI Apo LWD 40× WI 1.15 NA Lambda S and CFI Apo 20× WI 0.95 NA Lambda S. The laser lines used on the Zeiss microscopes were 488, 561, 568, 633, and 647 nm. A Nikon LUNV solid state laser launch was used for lasers 395/405, 488, 561 and 647 nm for GFP/Alexa488, RFP/Alexa546, and Alexa647 respectively. Emission filters used on the Nikon were 480/30, 535/30, 605/52. Nikon Elements Advanced Research v5.41.02 (Nikon) was used for image acquisition.

### Lineage tracing

*Tg(krtt1c19e:lyn-Tomato)^sq16^*and *Tg(dld:hist2h2l-EGFP)^psi84^*3–5 dpf larvae were anesthetized with buffered MS222 up to 150 mg/L and mounted in glass-bottomed dishes (Cellvis) using 0.8% low melting point agarose dissolved in DI water supplemented with buffered MS222 (120 mg/L). Time lapses were acquired in a Nikon Ti2 Yokogawa CSU-W1 spinning disk head equipped with a Hamamatsu Orca Fusion sCMOS and CFI Apo LWD 40× WI 1.15 NA Lambda S objective. A Stage Top Incubator (OkoLab) was used to keep the constant temperature of 28.5°C and 85% humidity.

### Serial-section electron microscopy (ssEM)

At 7 dpf, a larval zebrafish underwent aldehyde specimen fixation followed by heavy metal contrast staining for electron microscopy as described in (Petkova, 2020). The fish was embedded and cured in LX-112 resin, cut in 30 nm thick sections using an automated tape collection ultramicrotome (ATUM) system (Kasthuri et al., 2015) and mounted on silicon wafers (University Wafers, USA) for imaging. The image volume was obtained with a Zeiss multibeam scanning electron microscope (mSEM) equipped with 61 overlapping electron beams (Eberle et al., 2015) following the procedure outlined in (Shapson-Coe et al., 2024). Rigid stitching was performed on raw microscope tiles by extracting SIFT features, with global optimization applied to smooth the results across each section, and elastic stitching used for sections with distortions. Low-resolution thumbnails were generated in real-time, matched with nearby sections, and refined using a spring mesh model to prepare the images for final alignment. We used SOFIMA (Januszewski et al., 2024) to obtain a precise alignment of the complete stack, following the procedures outlined previously (Shapson-Coe et al., 2024). Briefly, starting from roughly prealigned sections we computed an optical flow field between each pair of adjacent sections using 8x8 nm^2^ EM images. The field vectors were estimated on a regular grid with 40 px spacing. We then modeled each section as an elastic spring mesh grid with edge sizes corresponding to the flow field vector spacing, and with additional 0-length springs representing the flow field vectors. We allowed the system to relax, regularizing the flow field and finding a solution balancing deformation of the original images and cross-sections alignment. We used the final state of the mesh to warp the images into alignment to obtain the 3D stack.

Sections were painted manually using VAST *Lite* version 1.4.1 (Berger et al., 2018). 3D objects were exported as mesh files using the ‘vasttools.m’ MATLAB script in VAST *Lite*, and processed in Fiji (Schindelin et al., 2012), using the 3D viewer plugin (Schmid et al., 2010). Empty sections in the dataset (marking the position of knife changes) were removed in Blender (blender.org).

### Scanning electron microscopy (SEM)

Zebrafish larvae (*ift88^-/-^* homozygous mutants, lacking cilia) raised in 1× E3 medium were fixed at 4 dpf in 2.5% glutaraldehyde/0.1M sodium cacodylate buffer overnight. Samples were washed in buffer, post-fixed in 2% aqueous osmium tetroxide for 1 hour, and washed in buffer again. Samples were dehydrated through a graded ethanol series (50, 75, 95, 100%), followed by 50:50 hexamethyldisilazane (HMDS):ethanol, and then 100% HMDS. After removal of the HDMS, samples were dried in a fume hood overnight. Samples were mounted onto a pin stub using a

Leit-C sticky tab and mounting putty, and gold-coated using an Edwards S150B sputter coater. Samples were imaged in a Tescan Vega3 LMU Scanning Electron Microscope (operating voltage, 15 kV) using a secondary electron detector.

### Image processing, quantifications, and statistical analyses

Zeiss LSM 800 confocal images were subjected to Gaussian Blur 3D processing (X:0, Y:0, Z:2) in Fiji. Zeiss LSM 880 Airyscan and LSM 980 Airyscan2 confocal images were subjected to Airyscan processing on Zen Blue 3.7 (Zeiss) using “Auto” Airyscan processing parameters. Further processing (for example, gamma correction, maximum intensity projections, and 3D rendering) was performed in Fiji. For fluorescence intensity quantification, background subtraction was performed with rolling ball radius of 50 pixels, and a ROI was drawn around each cell of interest. Quantification was performed using the Analyze, Measure function in Fiji. No image quantification was performed in gamma corrected images. Single channel confocal images may be presented with their grayscale values inverted. Bleed-through in HCR RNA-FISH imaging for Alexa488- and Alexa546-conjugated probes was removed in Fiji, by subtracting the calculated value.

Statistical analyses were carried out in GraphPad Prism 10. Datasets were considered normally distributed if they passed the Kolmogorov-Smirnov test. Subsequent statistical tests used are stated in the figure legends. Bars on graphs indicate mean ± standard error of the mean (S.E.M.).

Figures were prepared using Adobe Photoshop versions 25.9.0 and 25.11.0, Adobe Illustrator versions 25.4.1, 28.6, and 29.4, and Affinity Designer.

### Ethics Statement

This study was conducted in accordance with the Guide of the Care and Use of Laboratory Animals of the National Institutes of Health and the Singapore National Advisory Committee for Laboratory Animal Research. Protocols involving live animals were approved by the Institutional Animal Care and Use Committees of the Stowers Institute for Medical Research (TP Protocol: # 2023-159), Nanyang Technological University (A23136) and Harvard University (25-03).

## Results

### Single-cell RNA sequencing reveals the presence of several classes of olfactory ionocytes

To search for potential ionocytes in the zebrafish olfactory epithelium, we used single-cell RNA sequencing (scRNA-seq) datasets generated from dissected adult olfactory rosettes (Kraus et al., 2022). We identified both major types of olfactory sensory neurons (OSNs; ciliated and microvillous, marked by *ompb* and *trpc2b*, respectively), supporting cells, neuronal progenitors and immune cell clusters (Fig. 1A; Fig. S1A–C). Additionally, we found a cluster containing the well-conserved pan-ionocyte markers, *foxi3a* and *foxi3b (Hsiao et al., 2007; Jänicke et al., 2007)*. Differential gene expression among clusters shows the expression of NaR ionocyte genes such as *trpv6*, *gcm2*, *fxyd11* (*si:dkey-33i11.4*) (Saito et al., 2010) and *atp1a1a.3* (Liao et al., 2009), and putative HR ionocyte markers, such as *ceacam1* (Fig. 1B–F; cluster 18 in Table S1). The cluster contains a high level of mitochondrial genes (Fig. S1D), consistent with the mitochondria-rich characteristic of ionocytes.

**Figure 1.**
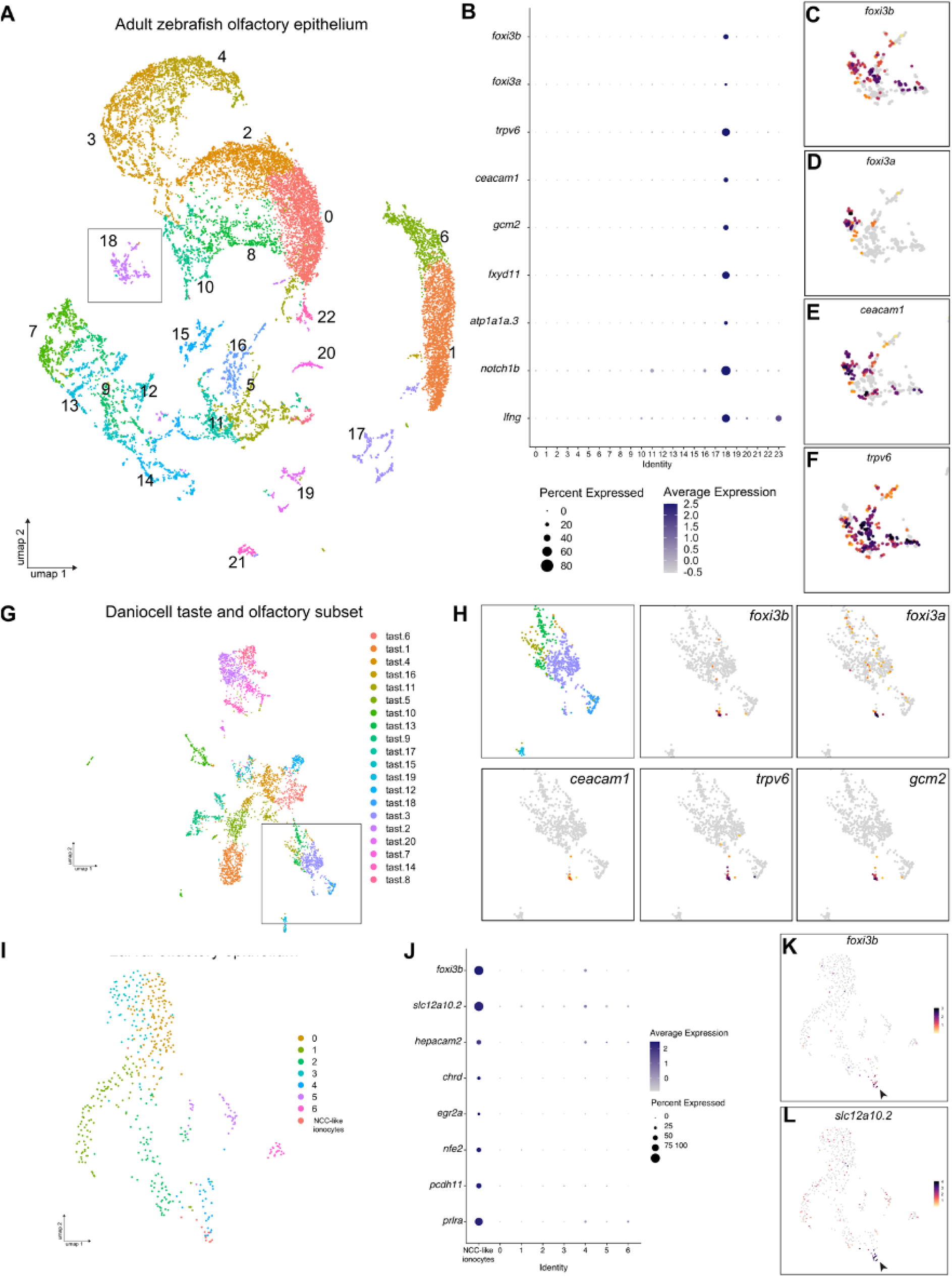
Single-cell RNA sequencing data analysis reveals expression of classical ionocyte-marker genes in the zebrafish olfactory organ. **(A)** UMAP plot showing 24 unannotated cell clusters (Identity) in the scRNA-seq dataset generated from dissected adult zebrafish olfactory organs. **(B)** Dot plot of data from A depicting several known ionocyte markers expressed in cluster 18. **(C)** Feature plots for *foxi3b*, (D) *foxi3a*, (E) *ceacam1*, and (F) *trpv6*. **(G)** UMAP plot of taste/olfactory subset from Daniocell, a scRNA-seq dataset generated from whole embryos and larvae (Sur et al., 2023). **(H)** Zoomed in feature plots for the ionocyte markers *foxi3b, foxi3a, ceacam1, trpv6*, and *gcm2*. **(I)** UMAP plot of olfactory subset from (Peloggia et al., 2021). **(J)** Dot plot of data from J showing several NCC ionocyte markers in *foxi3b^+^* cells in the olfactory epithelium. **(K)** Feature plots for *foxi3b* and (L) *slc12a10.2*.

We did not detect genes characteristic of other zebrafish ionocyte subtypes, such as *slc12a10.2* (Wang et al., 2009). Differential expression analysis also indicated the expression of genes that can contribute to Notch-Delta signalling (e.g. *lfng*, *notch1b*), similar to descriptions of different ionocyte subtypes in other tissues (Peloggia et al., 2021, 2024). These results suggest the presence of olfactory ionocytes in the adult zebrafish.

To determine if olfactory ionocytes are present at larval stages, we searched for the expression of ionocyte marker genes in scRNA-seq datasets from larval zebrafish (Farrell et al., 2018; Sur et al., 2023). In addition to the expression in previously defined ionocyte clusters, we found that the same genes were co-expressed in a small group of cells within the larval olfactory epithelium cluster (Fig. 1G,H). Interestingly, in a separate dataset from larval epithelial cells (Peloggia et al., 2021), we detected *foxi3b^+^* cells in the olfactory epithelium cluster that do not express genes that mark NaR and HR cells. Differential gene expression of these *foxi3b*^+^ cells against the remaining cells revealed a different set of ionocyte markers, including *slc12a10.2*, characteristic of NCC ionocytes (Fig. 1I–L; Table S2).

Transcriptomic data thus suggest the presence of three distinct subtypes of ionocytes in the larval olfactory epithelium and adult zebrafish olfactory rosette. These subtypes have transcriptional signatures of skin NaR, HR and NCC ionocytes.

### Spatial expression analysis identifies paired and solitary olfactory ionocytes

To validate the transcriptomic data, and to determine the localization and number of olfactory ionocytes in larval fish, we performed whole-mount fluorescent *in situ* hybridization with hybridization chain reaction (HCR RNA-FISH) (Choi et al., 2018) (Fig. 2; see Fig. 2A for anatomical location of olfactory pits in a DAPI-stained 5 dpf larva). We initially examined the expression of *foxi3b*, expressed in all ionocyte subtypes detected in the olfactory transcriptomes, and *trpv6* and *ceacam1* to label the NaR and HR ionocyte populations, respectively. *foxi3b*^+^ cells were found in all regions of the olfactory pit at 5 dpf, but were enriched in the posterolateral region (see white arrowhead; Fig. 2B–B’). There was a mean of 13 *foxi3b^+^* cells per olfactory pit (*N* of olfactory pits = 6).

**Figure 2.**
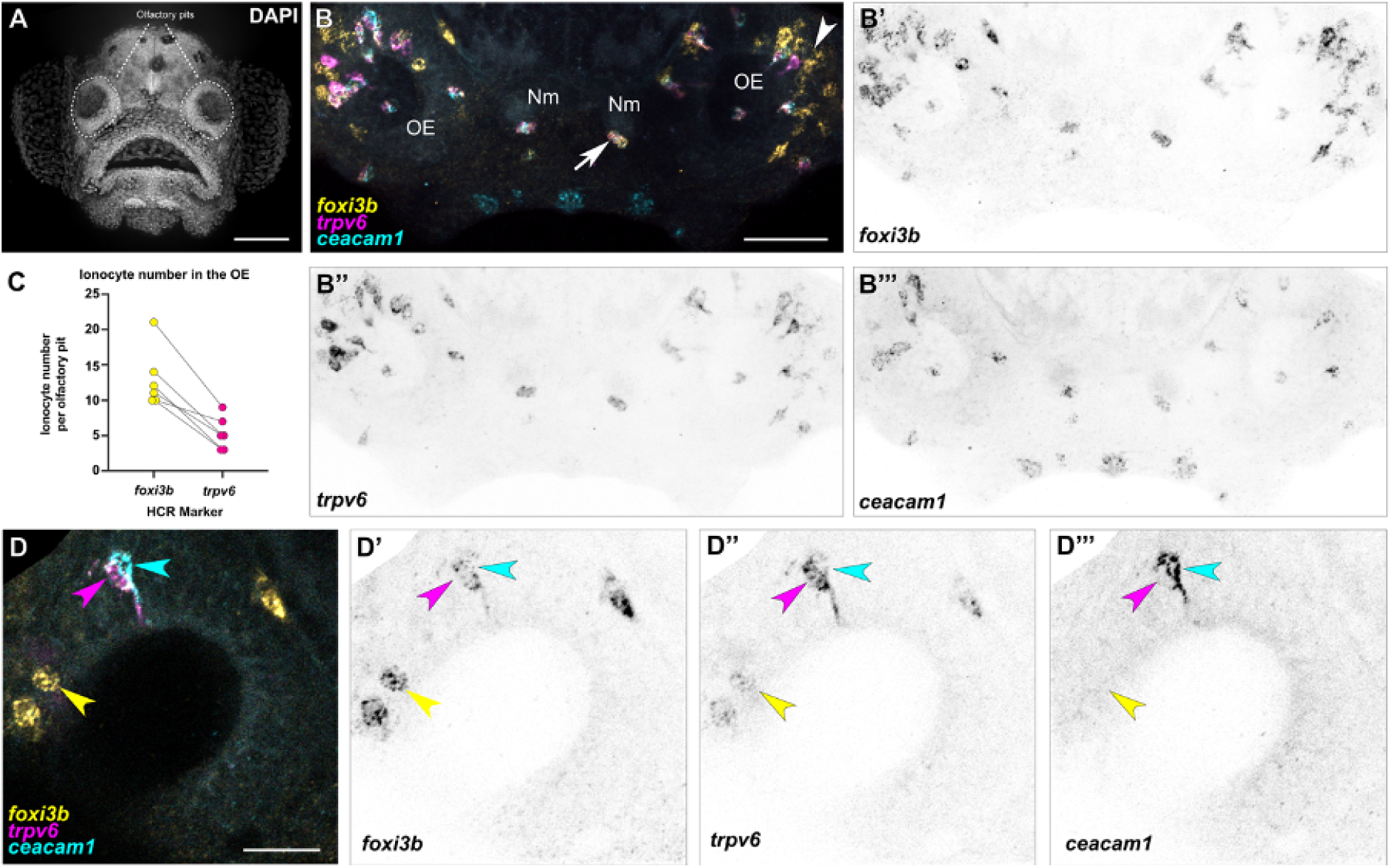
The larval zebrafish olfactory epithelium contains three distinct subtypes of ionocytes. (A) Maximum intensity projection of a 4 dpf larva stained with DAPI showing the two olfactory pits; frontal view. Scale bar: 50 µm. **(B–B’’’)** Maximum intensity projection of a confocal image of HCR RNA-FISH for *foxi3b* **(B’)**, *trpv6* **(B’’)**, *ceacam1* **(B’’’)**, and merged signals **(B)** in the head of a 5 dpf wild-type larva; dorsal view, anterior to the bottom. White arrowhead marks an example of an olfactory ionocyte in the posterolateral region of the olfactory pit with expression of *foxi3b*. White arrow marks an example neuromast ionocyte with expression of all three selected genes. Scale bar: 50 µm. Abbreviations: OE; olfactory epithelium, Nm; neuromast. **(C)** Numbers of *foxi3b*^+^ ionocytes per olfactory pit in 5 dpf larvae raised in 0.5× E2 medium, of which also express *trpv6*. Connecting lines indicate the same olfactory pit. **(D–D’’’)** Confocal image of HCR RNA-FISH signals for *foxi3b* **(D’)**, *trpv6* **(D’’)**, *ceacam1* **(D’’’)**, and merged signals **(D)** in the olfactory epithelium of a 5 dpf wild-type larva; dorsal view, anterior to the bottom, lateral to the left. Magenta and cyan arrowheads mark an example pair of ionocytes, with the magenta arrowhead marking a strong *trpv6*-expressing cell, and the cyan arrowhead marking a *ceacam1*^+^ cell with weak *foxi3b* expression. Yellow arrowhead marks an example solitary ionocyte, which has strong expression of *foxi3b* and weak expression of *trpv6*. Scale bar: 20 µm.

A subset of the *foxi3b^+^* cells expressed the NaR marker *trpv6* (Fig. 2C). Interestingly, we always detected one cell with strong *ceacam1* expression (HR-like) adjacent to a *trpv6*^+^ (NaR-like) cell (Fig. 2B–B’’’,D–D’’’). These pairs of cells typically had nuclei situated deep in the epithelium, and each had an extension reaching the epithelial surface (see magenta and cyan arrowheads, Fig. 2D; Movie S1).

Notch pathway genes are expressed in olfactory ionocytes (Fig. 1B). To test whether Notch signalling is active in these pairs, we used the Notch reporter *Tg(tp1bglobin:EGFP)^um14^*in combination with HCR RNA-FISH. Notch signalling is active in one of the two cells of the ionocyte pairs in the olfactory epithelium, with the NaR-like ionocyte (*trpv6^+^*) being the Notch-positive cell of the pair (Fig. 3A–A’’).

**Figure 3.**
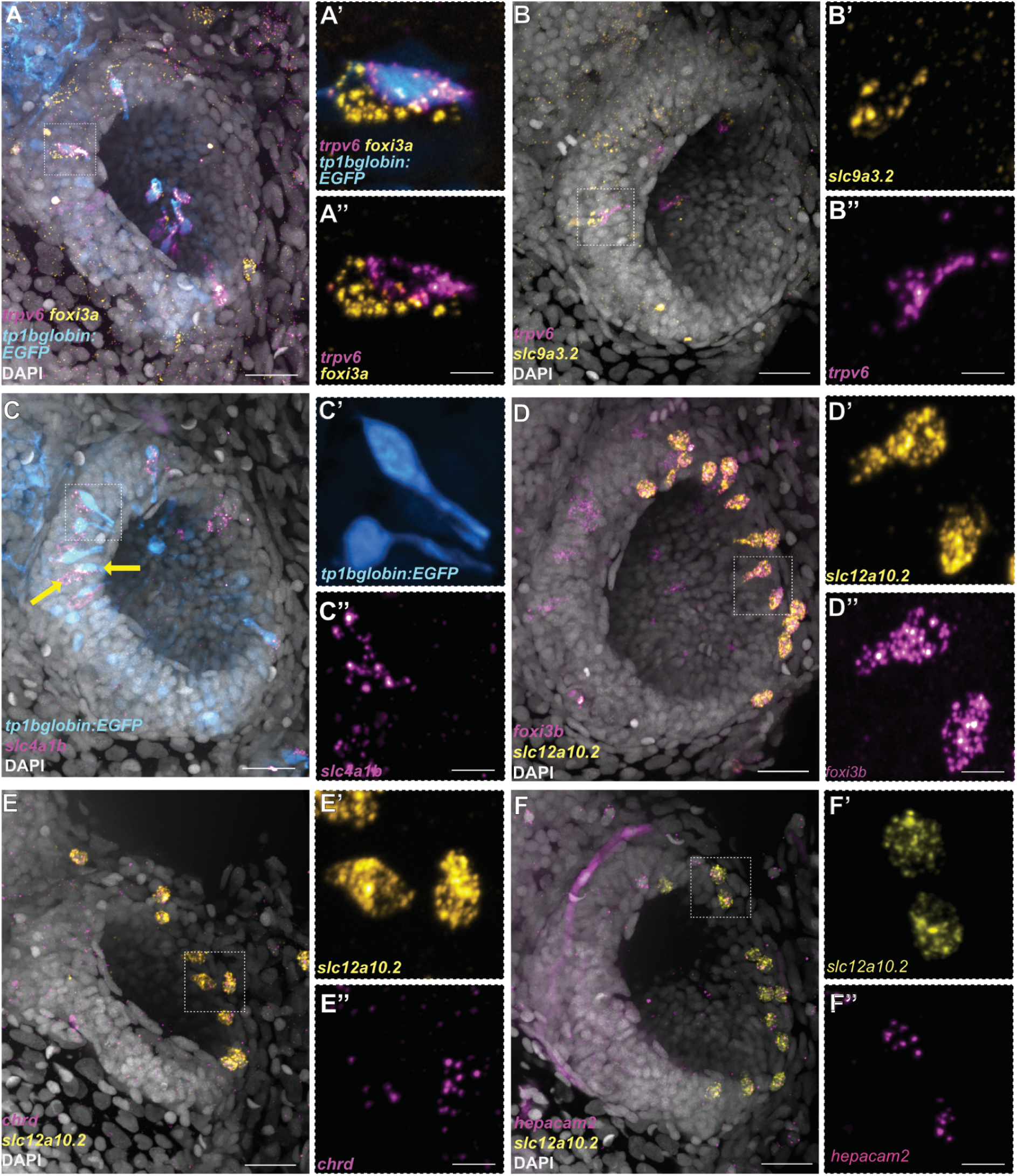
Differential expression of Notch reporter and ion channel genes in the three distinct subtypes of olfactory ionocytes. (A–A’’) HCR RNA-FISH for *foxi3a* (yellow) and *trpv6* (magenta), combined with the Notch reporter *tp1bglobin:EGFP* (cyan) and DAPI stain (grey). (B–B’’) HCR RNA-FISH for *slc9a3.2* (yellow) and *trpv6* (magenta) with DAPI stain (grey). (C–C’’) HCR RNA-FISH for *slc4a1b* (magenta) combined with the Notch reporter *tp1bglobin:EGFP* (cyan) and DAPI stain (grey). The arrows indicate adjacent cells with EGFP and *slc4a1b* expression. **(D–D’’)** Maximum intensity projection of a HCR RNA-FISH for *slc12a10.2* (yellow) and *foxi3b* (magenta) with DAPI stain (grey) shows solitary, NCC-like ionocytes in the olfactory epithelium. **(E–E’’)** HCR RNA-FISH for *slc12a10.2* (yellow) and *chrd* (magenta) with DAPI stain (grey). **(F–F’’)** HCR RNA-FISH for *slc12a10.2* (yellow) and *hepacam2* (magenta) with DAPI stain (grey). Scale bars: **A**, **B**, **C**, **D**, **E**, **F**, 20 µm; **A’’**, **B’’**, **C’’**, **D’’**, **E’’**, **F’’**, 5 µm.

To test which other ion channel genes are expressed in these cells, we performed additional stainings for HR markers. Indeed, the *trpv6^-^* cell of the pair expresses the sodium/proton exchanger *slc9a3.2* (Fig. 3B–B’’) and the anion exchanger *slc4a1b* (Fig. 3C–C’’). However, these cells do not express the ammonium transporter *rhcgb* (Fig. S2), which has been detected in HR ionocytes in the skin (Nakada et al., 2007).

Besides the presence of *trpv6^+^* and *ceacam1^+^*ionocyte pairs, we also observed solitary *foxi3b*^+^ cells that were not paired with *ceacam1*^+^ cells (Fig. 2D). These cells were positioned on the lateral borders of the olfactory pit and had a rounded or teardrop-shaped morphology (see yellow arrowhead, Fig. 2D; Movie S2). To test if these *foxi3b^+^* cells correspond to the NCC-like cells present in the larval epithelial dataset, we performed HCR RNA-FISH, combining a *foxi3b* probe with different markers obtained from differential expression analysis of scRNA-seq data. We confirmed that these solitary *foxi3b^+^*cells express the NCC ionocyte markers *slc12a10.2*, *chrd*, and *hepacam2* (Peloggia et al., 2021; Casellas et al., 2023) (Fig. 3D–F’’).

We used HCR RNA-FISH to examine the distribution of ionocytes in adult zebrafish olfactory rosettes. Ionocyte pairs were detected throughout the rosette (Fig. 4A–B’’’); *trpv6*^+^ cells were adjacent to a strongly *ceacam1*^+^ cell throughout the epithelium, with both cell types having varying levels of *foxi3b* expression (Fig. 4B’’’,C’’’,D’’’). In the peripheral non-sensory region, by contrast, we additionally observed *foxi3b^+^* cells with no expression of *trpv6* or *ceacam1* (Fig. 4D–E’’’). These NCC-like ionocytes appeared rounded in all cases (Fig. 4D,D’’’,E,E’’’; Fig. S3), whereas NaR-like/HR-like ionocytes had variable morphologies, appearing elongated in the region close to the OSN zone, and rounded in the most peripheral non-sensory regions (Fig. 4D–E’’’).

**Figure 4.**
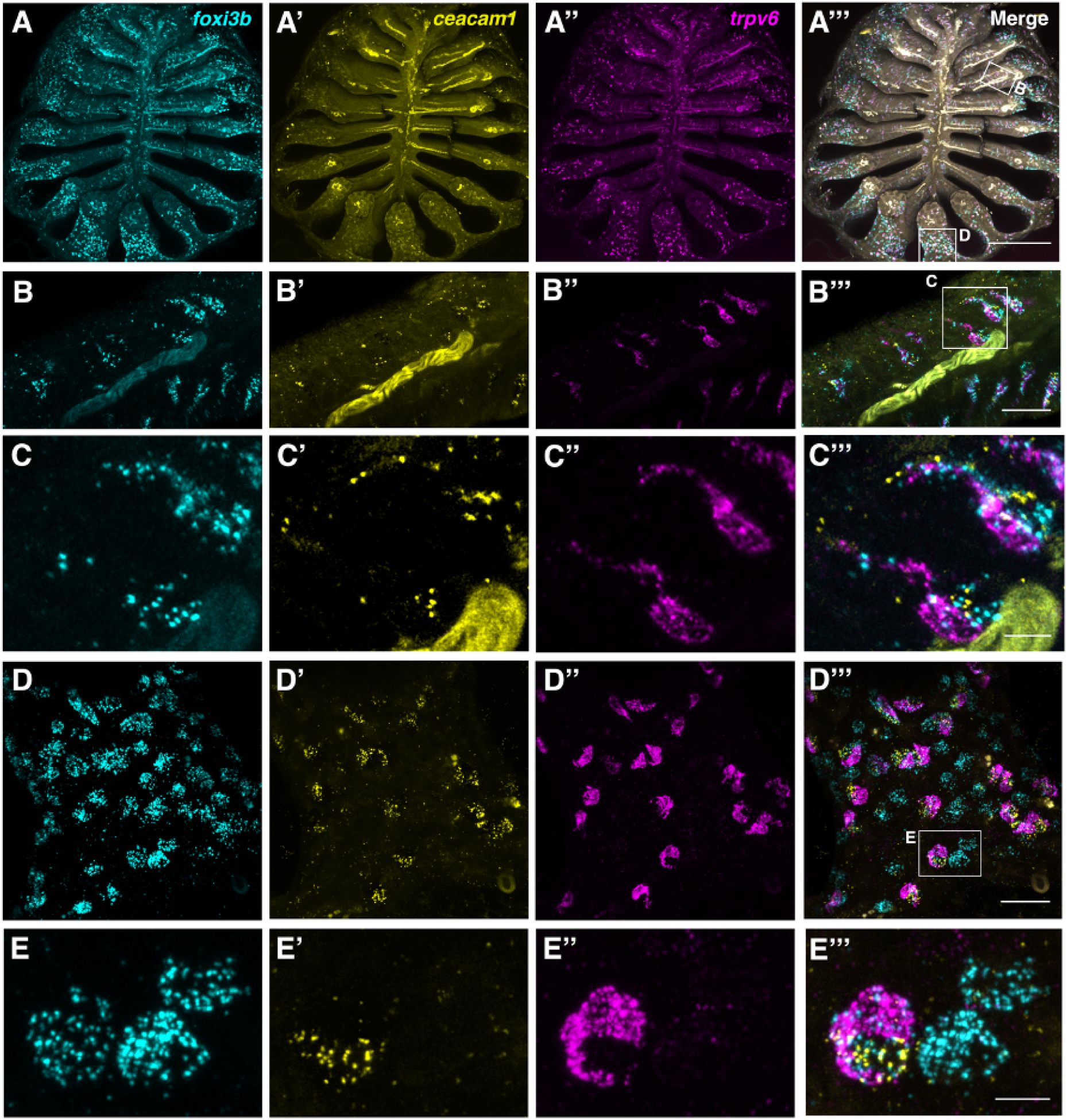
The adult zebrafish olfactory epithelium contains three distinct subtypes of ionocytes. (A–A’’’) Overview of a dissected adult olfactory rosette. Maximum intensity projections of an Airyscan2 confocal image of HCR RNA-FISH for *foxi3b* **(A)**, *ceacam1* **(A’)**, *trpv6* **(A’’)**, and merged signals **(A’’’)**. Scale bar: 200 µm. (**B–B’’’**) Enlargement of the boxed region ‘B’ in A’’’, within the central (sensory) zone of the olfactory rosette. Maximum intensity projections of a subset of optical sections; HCR RNA-FISH for *foxi3b* **(B)**, *ceacam1* **(B’)**, *trpv6* **(B’’)**, and merged signals **(B’’’)**. Pairs of elongated ionocytes with cell bodies located deep in the epithelium are visible. (The yellow stripe running through the image is autofluorescence from a blood vessel.) Scale bar: 20 µm. (**C–C’’’**) Enlargement (maximum intensity projection of a subset of *z*-slices used in B) of boxed region in B’’’, featuring two ionocyte pairs: HR-like ionocytes expressing *ceacam1* (yellow) and *foxi3b* (cyan), adjacent to NaR-like ionocytes expressing *trpv6* (magenta) and a low level of *ceacam1*. HCR RNA-FISH for *foxi3b* **(C)**, *ceacam1* **(C’)**, *trpv6* **(C’’)**, and merged signals **(C’’’)**. Scale bar: 5 µm. **(D–D’’’)** Enlargement of boxed region ‘D’ in A’’’, within the peripheral (non-sensory, multiciliated) zone of the olfactory rosette. Maximum intensity projections of a subset of *z*-slices used in A; HCR RNA-FISH for *foxi3b* **(D)**, *ceacam1* **(D’)**, *trpv6* **(D’’)**, and merged signals **(D’’’)**. Both paired and solitary ionocytes are present. Scale bar: 20 µm. **(E–E’’’).** Enlargement (maximum intensity projection of a subset of *z*-slices) of the boxed region in D’’’, featuring one HR-like/NaR-like ionocyte pair, and two NCC-like ionocytes. An HR-like ionocyte, expressing *ceacam1* (yellow) and *foxi3b* (cyan), sits adjacent to an NaR-like ionocyte expressing *trpv6* (magenta) and a lower level of *ceacam1*. The NCC-like ionocytes express *foxi3b* (cyan) but not *ceacam1* or *trpv6*. Ionocytes near the periphery of the rosette were rounded in shape. Scale bar: 5 µm.

We conclude that the zebrafish olfactory epithelium contains three main types of ionocytes, at both larval and adult stages. One type is an NCC-like ionocyte, which is solitary and expresses the sodium/chloride symporter *slc12a10.2*. The other two types, which are present in pairs, consist of one *trpv6^+^*, NaR-like ionocyte and one *ceacam1^+^*, HR-like ionocyte.

### Time-course of olfactory ionocyte development

To determine when olfactory ionocytes appear during embryonic development and their dynamics, we performed a time course analysis of *foxi3b* and *trpv6* expression from 1 to 5 dpf (Fig. 5). None of the ionocyte subtypes were observed in embryos at 1 dpf. Most olfactory pits showed solitary *foxi3b*^+^ ionocytes at 2 dpf, while *trpv6*^+^ ionocytes with an adjacent *foxi3b*^+^ cell were not as frequent (Fig. 5A,B). The numbers of all olfactory ionocyte subtypes progressively increased over time (Fig. 5A–C).

**Figure 5.**
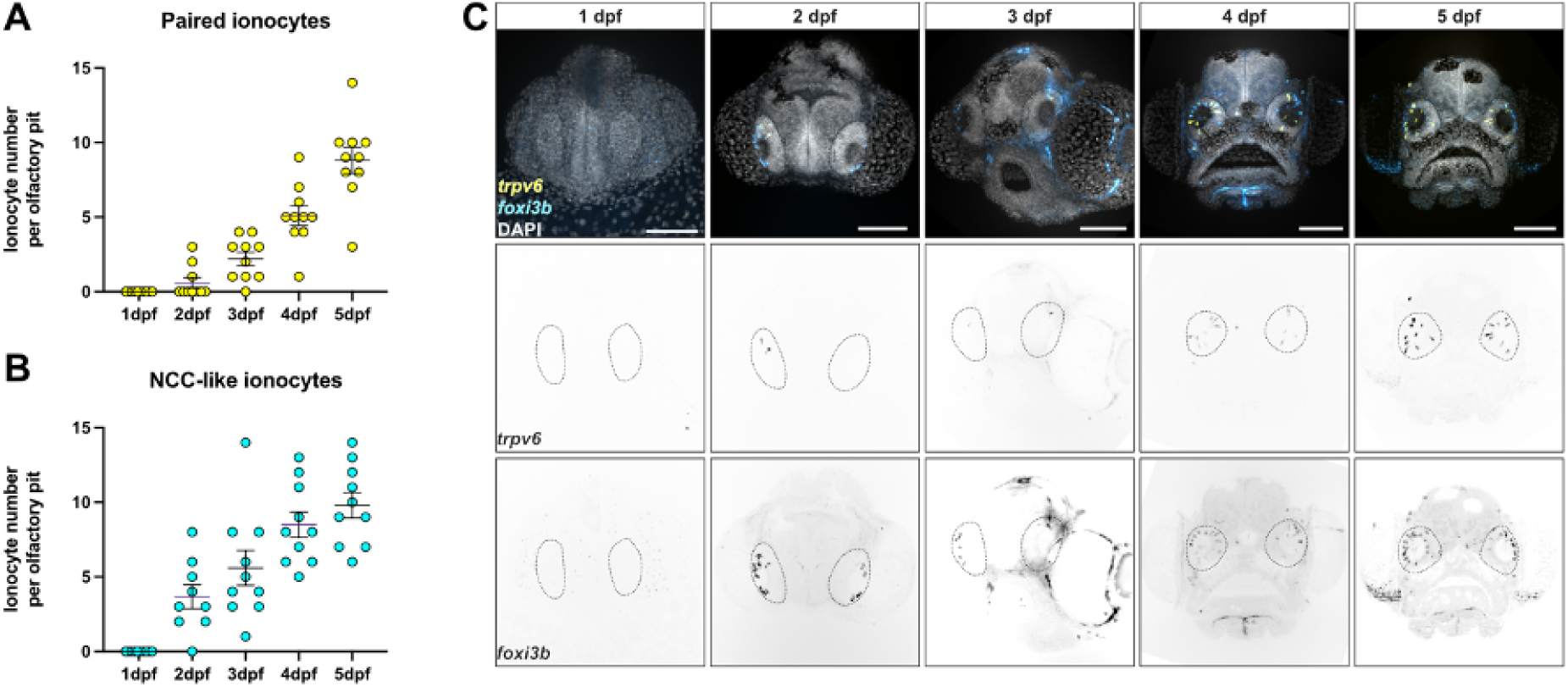
The subtypes of olfactory ionocytes develop at different times. **(A)** Developmental time course of NaR-/HR-like ionocyte pairs and **(B)** Number of NCC-like ionocytes from 1 to 5 dpf observed by confocal images of HCR RNA-FISH. **(C)** Representative maximum intensity projection confocal images of HCR RNA-FISH for *trpv6* (yellow) and *foxi3b* (cyan) with DAPI stain (grey) from A and B. Scale bar: 20 µm

Besides the presence of mature ionocytes, we also detected *foxi3a^+^*cells adjacent to the olfactory epithelium (Fig. S4A). These cells did not express any of the ion channel or transporter genes we examined by HCR RNA-FISH. Similar cells have been observed in the lateral line neuromast (Peloggia et al., 2021). In these organs, neuromast ionocyte progenitors are *krt1-19e^+^* skin cells that turn off expression of the ionocyte specification transcription factor genes *foxi3a* and *foxi3b* as they invade neuromasts, where they differentiate. To test if *krt1-19e^+^* cells give rise to new ionocytes in the olfactory epithelium during development, we performed time-lapse analyses of the transgenic line *Tg(krtt1c19e:lyn-tdTomato)^sq16^*. While we observed several new ionocytes invading the lateral line neuromasts, we did not observe any *tdTomato^+^* cells migrate into the olfactory pit (Fig. S4B,C; Movie S3). Additionally, olfactory pits did not contain any *tdTomato*^+^ cells, suggesting that these cells do not give rise to ionocytes in the olfactory epithelium (Fig. S4B,C’’). Thus, ionocytes in the olfactory epithelium have a distinct origin from those in neuromasts: they may originate outside the olfactory epithelium, but do not appear to derive from *krt1-19e^+^* cells.

### Notch signalling differentially regulates olfactory ionocyte number

Ionocyte differentiation and survival is regulated by Notch signalling in different tissues (Hsiao et al., 2007; Jänicke et al., 2007; Peloggia et al., 2021, 2024). To test if Notch signalling also plays a role in ionocyte development and maintenance in the olfactory epithelium, we treated 4 dpf larvae for 24 hours with the gamma-secretase inhibitor LY411575, which inhibits Notch signalling (Fig. 6A). We observed a striking loss of elongated *trpv6*^+^ and adjacent *foxi3b*^+^ cells, which are likely the NaR- and HR-like pairs, but no change to solitary, posterolaterally located *foxi3b*^+^ cells, which are likely NCC-like ionocytes (Fig. 6B–E’’). The remaining NaR- and HR-like pairs show altered cell morphology, and could be undergoing cell death (Fig. 6E–E’’).

**Figure 6.**
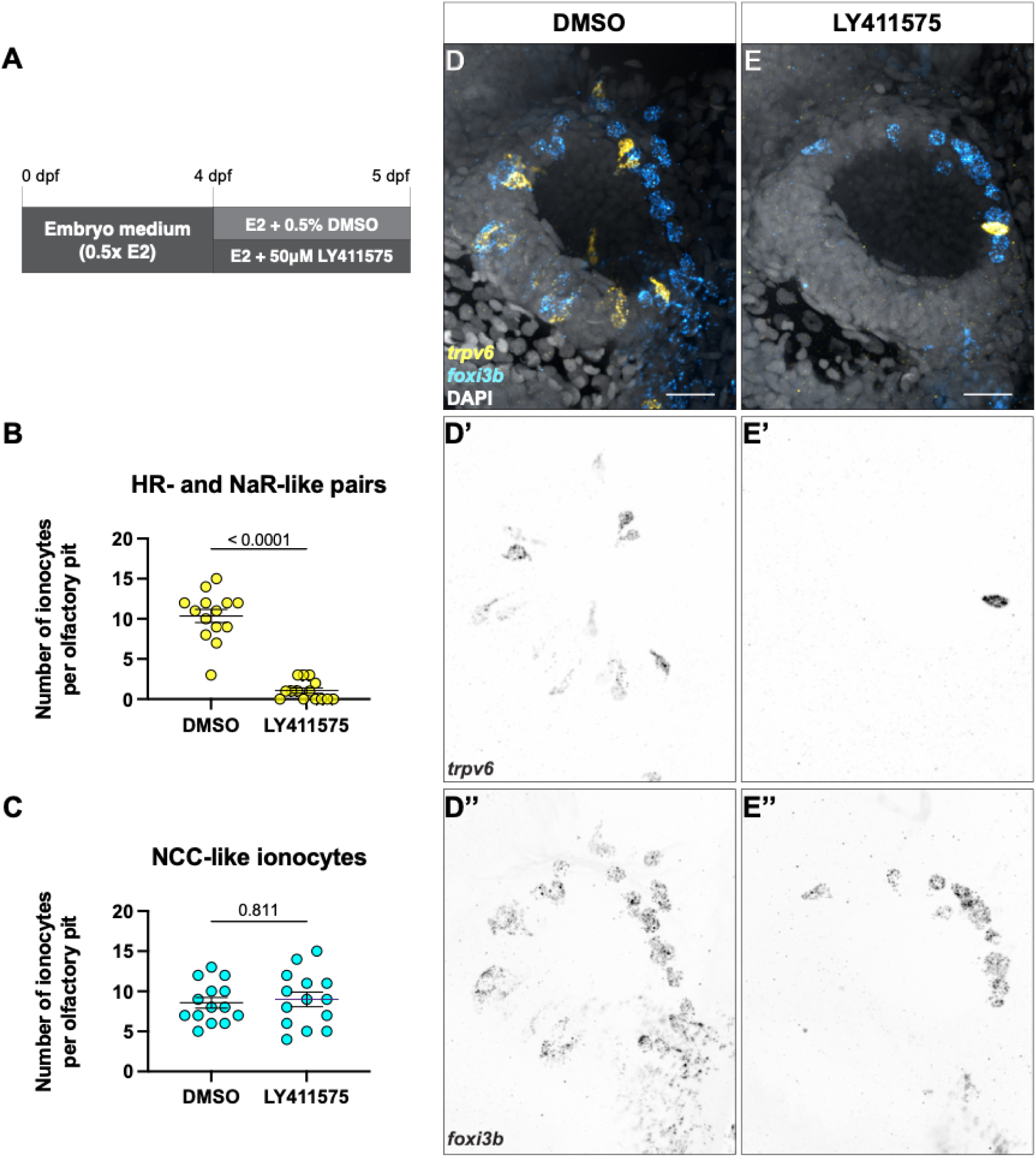
Notch signalling is required for paired ionocyte survival, but does not affect NCC-like ionocyte number. **(A)** Schematic of experimental design. **(B)** Number of NaR- and HR-like ionocyte pairs in DMSO controls and treated with the Notch inhibitor LY411575. Mann Whitney test (*P* < 0.0001). **(C)** Number of NCC-like ionocytes in DMSO controls and treated with the Notch inhibitor LY411575. Mann-Whitney test (*P =* 0.7811). **(D–E’’)** Representative maximum intensity projection confocal images of HCR RNA-FISH for *trpv6* (yellow) and *foxi3b* (cyan) with DAPI stain (grey) in DMSO-treated **(D–D’’)** and LY411575-treated **(E–E’’)** olfactory pits. *N =* 14 olfactory pits per condition. Scale bars: 20 µm.

In neuromast ionocytes, transcription of the Notch ligand gene *dld* is upregulated during division of the ionocyte progenitor cell into the NaR- and HR-like pair, resulting in two *dld^+^* cells (Peloggia et al., 2021). To investigate whether olfactory NaR- and HR-like pairs develop in a similar manner, we performed time-lapse analysis of the transgenic *dld* reporter line *Tg(dld:hist2h2l-EGFP)^psi84Tg^* from 3–5 dpf. We observed pairs of EGFP^+^ cells appear at the edge of the pit, upregulate EGFP and differentiate into ionocytes (Movies S4, S5). These data suggest that, similar to neuromast ionocytes, NaR- and HR-like ionocyte pairs in the olfactory epithelium do not originate from the division of pre-existing ionocyte pairs, but come from a different population of progenitor cells.

### Ultrastructure of larval zebrafish olfactory ionocytes

To describe the ultrastructure and three-dimensional shape of olfactory ionocytes, we examined a serial-section electron microscopy (ssEM) dataset of a 7 dpf wild-type zebrafish larva. Here, we found mitochondria-rich cells in the olfactory epithelium with ultrastructural features typical of teleost ionocytes (Bertmar, 1972; Karnaky et al., 1976; Pisam et al., 1983; Ruzhinskaya et al., 2001; Laurent et al., 2006; Fridman, 2020). An extensive tubular network gave the cytoplasm a lacy appearance, quite distinct from that of OSNs or other olfactory cell types, making it possible to spot these relatively rare cell types. Consistent with the HCR RNA-FISH data, the cells had differing shape and appearance in different regions of the olfactory epithelium.

We found several examples of ionocytes within the OSN zone of the olfactory pit (Fig. 7). These slender cells spanned almost the full width of the epithelium (>20 µm in apicobasal length), with nuclei positioned just above basal cells near the basal lamina. In line with the transcriptomics data, most examples consisted of a pair of cells. One member of the pair, which we propose is the HR-like cell (see below), terminated in an apical knob bearing ∼50 short, irregular microvilli, intermediate in diameter (0.2 µm) between the microvilli of microvillous OSNs (0.1 µm diameter) and the cilia of ciliated OSNs and non-sensory multiciliated cells (MCCs; 0.25–0.3 µm diameter) (Fig. 7A,C,E,E’; Movie S6). Apart from a cortical zone at the cell apex, the cytoplasm was densely packed with mitochondria in close association with an extensive intracellular tubular network (Fig. 7C,D).

**Figure 7.**
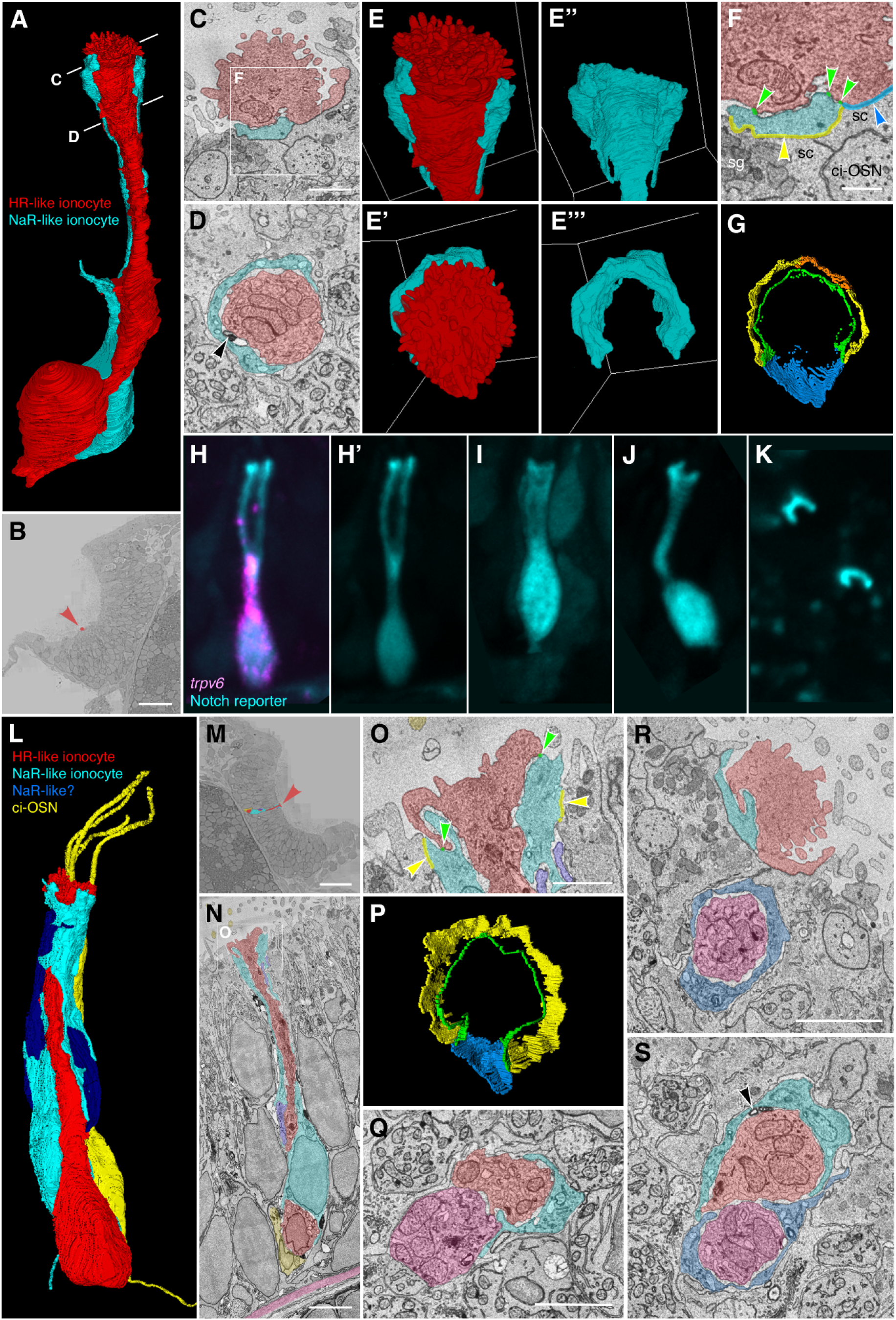
Ultrastructure and 3D reconstruction of olfactory HR-like/NaR-like ionocyte cell pairs and multicellular complexes in the wild-type zebrafish larva. (A–G) Representative example of an ionocyte cell pair in the 7 dpf zebrafish larval olfactory pit. The HR-like ionocyte is shown in red, with the NaR-like ionocyte in cyan. **(A)** Volume-EM 3D reconstruction of the cell pair; see also Supplementary Movie 6. **(B)** Location of the cell pair (red arrowhead) in the left olfactory pit (just within the OSN zone). Coronal section; anterior to the top. **(C,D)** Sections at the approximate levels shown in A. Black arrowhead in D marks an electron-dense structure in the extracellular space between the cell pair (see also S). **(E–E’’’)** 3D reconstructions of the apical part of the cell pair, showing the microvillous apical knob of the HR-like ionocyte (red), which projects above the surrounding olfactory supporting cells. Side views (top panels) and top-down views (lower panels). The neck of the HR-like ionocyte is wrapped by a thin layer of cytoplasm of the NaR-like cell (cyan). **(F,G)** Tight junctions (zonulae occludentes) of the ionocyte pair. **(F)** Enlargement of the box in C, showing color-coded labelling of the junctions. Green, shallow tight junction between the two ionocytes; yellow, deep tight junction between NaR-like ionocyte and olfactory supporting cell; orange, deep tight junction between NaR-like ionocyte and multiciliated cell; blue, deep tight junction between HR-like ionocyte and olfactory supporting cell. **Abbreviations:** ci-OSN, ciliated olfactory sensory neuron; sc, olfactory supporting cell; sg, secretory granule. **(G)** 3D reconstruction of the tight junctions (top-down view); see also Supplementary Movie 7. **(H–K)** Examples of NaR-like olfactory ionocytes in a live 5 dpf embryo, labelled by EGFP (cyan) in the Notch reporter line *Tg(tp1bglob:EGFP)*. **(H)** Co-expression of EGFP (Notch reporter; cyan) with HCR RNA-FISH for *trpv6* (magenta), confirming the cell as an NaR-like ionocyte (see also Fig. 3B–B’’). **(H’,I–K)** EGFP (cyan) channel only. **(I,J)** Additional examples in longitudinal view. **(K)** The apices of EGFP^+^ cells (NaR-like ionocytes) appear as crescents in a top-down view. **(L–S)** Examples of three- and four-cell ionocyte complexes in the wild-type zebrafish olfactory epithelium at 7 dpf. **(L–P)** Example of a 3-cell complex, consisting of an HR-like ionocyte (red), NaR-like ionocyte (cyan), and possible second NaR-like ionocyte (dark blue). A nearby ciliated OSN (yellow) is included for context. **(L)** 3D reconstruction of the ionocyte complex and ciliated OSN; see also Supplementary Movie 8. **(M)** Location of the cells in the OSN zone of the right-hand olfactory pit (red arrowhead). Coronal section; anterior to the top. **(N)** Longitudinal section through the complex, showing close association between the HR-like ionocyte (red) and the ciliated OSN (yellow) at the base, and location relative to the basal lamina (pink). **(O)** Enlargement of the box in N, showing color-coded labelling of the tight junctions of the HR-like (red) and NaR-like (cyan) cells (color code as in F). **(P)** 3D reconstruction of the tight junctions of the HR-like (red) and NaR-like (cyan) cells (top-down view; compare to G; see also Supplementary Movie 9). **(Q)** Example section through a three-cell complex consisting of an HR-like cell (red), an NaR-like cell (cyan), and possible second HR-like cell (pink). **(R,S)** Example sections through a four-cell complex consisting of two HR-like/NaR-like pairs. The cell pairs are separate at their apices (R), but are closely associated beneath the epithelial surface (S). Black arrowhead in S marks electron-dense structures between the ionocytes. **Scale bars: B**, 20 µm; **C**; 1 µm (applies to **D–E’’’**, **G**); **F**, 0.5 µm; **M**, 20 µm; **N**, 3 µm; **O**, 1 µm (applies to **P**); **Q**, 2 µm; **R**, 2 µm (applies to **S**).

The HR-like ionocytes within the OSN zone were closely associated with a second cell along most of their apicobasal length (Fig. 7). This second cell wrapped around the HR-like ionocyte at the cell apex with a thin layer of cytoplasm, forming a crescent in transverse section (Fig. 7A–E’’’). At their apices, the two cells were connected with a continuous shallow (0.1–0.2 µm) tight junction (zonula occludens), and to surrounding cells with deep (0.5–1 µm) tight junctions (Fig. 7F,G,L). In a gap not covered by the wrapping cell, the ionocyte was sealed to an olfactory supporting cell by a deep tight junction (Fig. 7F,G; Movie S7). The cytoplasm of the wrapping cell also had some ionocyte-like characteristics (e.g. some tubules and pores). The ultrastructure of both cells of the pair was clearly distinct from the supporting cells that surround and insulate the OSNs, which are full of secretory granules (Fig. 7C).

To determine which member of the cell pair corresponded to which type of ionocyte, we compared the morphology of the ssEM 3D reconstructions to that of live cells imaged at 5 dpf with the *Tg(tp1bglobin:EGFP)* Notch reporter line, which marks the NaR-like ionocytes (Fig. 7H–K). The morphology of the live NaR-like cells clearly matched that of the wrapping cells seen in the ssEM dataset, with a thin curved layer of cytoplasm near the cell apex, sometimes appearing as a doublet in a lateral view (Fig. 7H,H’), and forming a clear crescent in a top-down view (Fig. 7K). Taken together, the ssEM and fluorescence imaging data strongly suggest that the ionocyte cell pairs found in the ssEM dataset correspond to the HR-like and NaR-like ionocyte pairs identified through transcriptomic profiling, with the NaR-like cell wrapping around the apex of the HR-like cell.

We also found occasional examples of multicellular ionocyte complexes consisting of three or four cells within the OSN zone of the larval olfactory epithelium (Fig. 7L–S; Movies S8, S9). In one example (Fig. 7L–P), one HR-like/NaR-like pair was associated with a second possible NaR-like cell. In this example, the HR-like ionocyte was closely associated with a ciliated OSN at its base. In another example (Fig. 7Q), the cell complex appeared to consist of an HR-like/NaR-like pair with a second possible HR-like cell. We also found a four-cell complex consisting of a pair of HR-like/NaR-like pairs, separate at their apices, but closely associated beneath the surface of the epithelium (Fig. 7R,S).

Olfactory ionocytes with a different morphology were present in the multiciliated cell zone at the periphery of the olfactory pit (Fig. 8). Here, the rounded microvillous apical knobs of individual ionocytes could be identified by scanning electron microscopy (SEM) in *ift88^-/-^* mutant larvae at 4 dpf (Fig. 8A–C). These mutants lack cilia, which would otherwise obscure these cells in the wild-type olfactory pit. Ionocytes were visible in the olfactory pits of two out of three individuals. In the ssEM dataset of a wild-type larva at 7 dpf, mitochondria-rich cells containing a dense tubular network were present within the non-sensory zone of multiciliated cells (Fig. 8D–M). Some of these peripheral ionocytes were cuboidal in shape, without the long narrow neck of the ionocytes in the central OSN zone, where the olfactory epithelium is thicker (Fig. 8D–I). Other ionocytes in the multiciliated cell zone were more columnar in shape (Fig. 8J). Peripheral ionocytes formed deep tight junctions with surrounding non-sensory multiciliated cells, skin cells, and other ionocytes.

**Figure 8.**
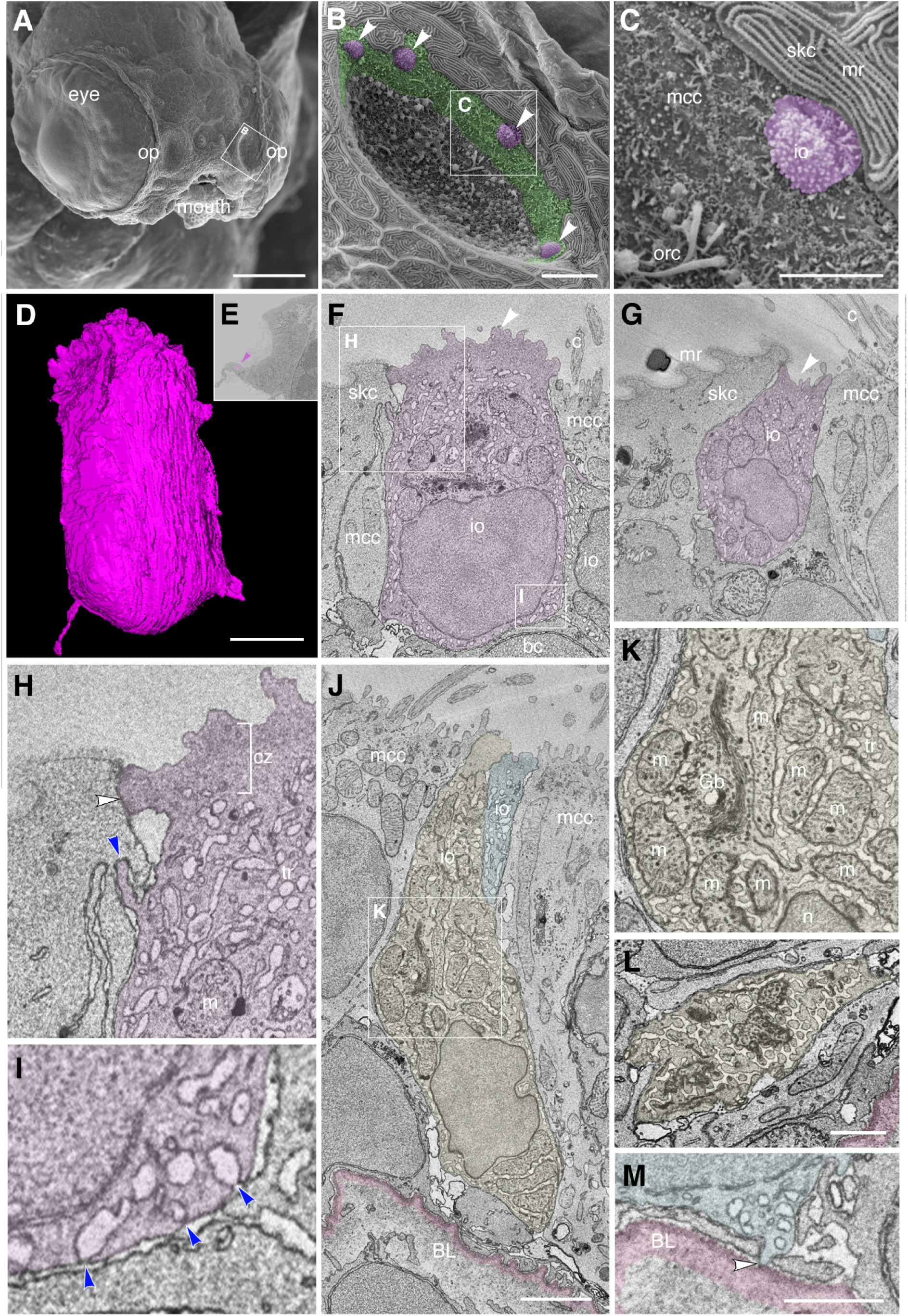
Ultrastructure and 3D reconstruction of NCC-like olfactory ionocytes in the non-sensory multiciliated cell zone of the zebrafish olfactory pit. (A–C) Scanning electron micrographs of an *ift88^-/-^* zebrafish mutant embryo at 4 dpf. **(A)** Whole head showing location of the two olfactory pits (op). **(B)** Enlargement of the left-hand olfactory pit, boxed in A. The rounded apical surfaces of four ionocytes (presumed NCC-like) are highlighted in magenta (arrowheads). The peripheral zone of non-sensory multiciliated cells (mcc) is highlighted in green. (All olfactory cilia are missing in the *ift88^-/-^* mutant, allowing visualization of the apical surface of cells in the pit.) **(C)** Enlargement of the boxed region in B. An ionocyte (magenta) sits in the MCC zone, in contact with a skin cell with microridges (mr, top right). The rods of two or three olfactory rod cells are also visible (orc, bottom left; see also Fig. 3I in (Cheung et al., 2021)). **(D–M)** Ultrastructure and 3D reconstruction of ionocytes in the non-sensory multiciliated zone of the olfactory pit in a wild-type zebrafish larva at 7 dpf. **(D)** 3D reconstruction of a presumed NCC-like ionocyte, showing the microvillous apical surface. **(E)** Location of the ionocyte in D (magenta; arrowhead) at the lateral edge of the left olfactory pit. Coronal section; anterior to the top. **(F,G)** Selected sections through the ionocyte shown in D, highlighted in magenta. The ionocyte makes contact with at least four other cell types: apically, with a skin cell on one side and a multiciliated cell on the other; basolaterally, with multiciliated cells, a basal cell, and another ionocyte. The microvillous apical surface is rounded in one area (F, arrowhead) but also forms a pit-like structure (G, arrowhead) in the same cell. **(H)** Enlargement of the boxed region in F (top left), highlighting tight junctional contact (white arrowhead) and interdigitation (blue arrowhead) between the ionocyte and a neighbouring skin cell. **(I)** Enlargement of the boxed region in F (bottom right), showing that pores where the tubular reticulum meets the plasma membrane are covered by a thin electron-dense structure (blue arrowheads). **(J)** Two examples of more elongated ionocytes highlighted in yellow and blue, with their apices sitting between multiciliated cells. **(K)** Enlargement of the boxed region in J, showing the mitochondria-rich cytoplasm, Golgi apparatus and extensive tubular reticulum. **(L)** Section through the base of the yellow cell in J, showing the tubular reticulum. **(M)** An end-foot (arrowhead) of the blue ionocyte in J makes direct contact with the basal lamina (pink). **Abbreviations:** bc, basal cell; BL, basal lamina (pink); c, cilia of multiciliated cell; cz, cortical zone (free of mitochondria; bracketed in H); Gb, Golgi body; io, presumed NCC-like ionocyte; m, mitochondrion; mcc, multiciliated cell; mr, microridges on skin cell; n, cell nucleus; op, olfactory pit; orc, olfactory rod cell (apical rods visible); skc, skin cell; tr, tubular reticulum. **Scale bars: A**, 100 µm; **B**, 10 µm; **C**, 5 µm; **D**, 2 µm (applies to **F, G**); **J**; 2 µm; **L**, 1 µm; **M**, 1 µm.

Basally, they were positioned just above the basal cells of the olfactory epithelium (Fig. 8F,J), occasionally extending thin end-feet to contact the basal lamina directly (Fig. 8M). However, they did not appear to be paired with, or wrapped by, any other cell along their entire apicobasal length, in contrast to the ionocytes in the OSN zone. Some, but not all, peripheral ionocytes had a large apical knob bearing ∼100 irregular microvilli, with part protruding from the epithelium (Fig. 8D,F), and part sunken to form a crypt or pit (Fig. 8G), matching the morphology revealed by SEM at 4 dpf (Fig. 8C). Based on their location in the multiciliated cell zone and morphology, and in comparison to the scRNA-seq data, we identify the cuboidal peripheral olfactory ionocytes as NCC-like cells.

## Discussion

Through analysis of single-cell transcriptomic, gene expression and volume electron microscopy data, we have identified at least three subtypes of ionocytes in the larval and adult zebrafish olfactory epithelium (Figs. 9, 10). All three express the pan-ionocyte marker *foxi3b*, but can be distinguished by transcriptomic signatures and morphology (Figs. (9A, 10A). NaR-like ionocytes, expressing *trpv6*, were paired with HR-like ionocytes, expressing *ceacam1* and the transporter genes *slc9a3.2* and *slc4a1b*. NaR-like/HR-like cell pairs were situated throughout the olfactory epithelium, including within sensory regions, with cell nuclei deep in the epithelium and a long protrusion extending to the epithelial surface. NCC-like ionocytes, expressing *slc12a10.2*, *chrd*, and *hepacam2*, were located exclusively in the multiciliated cell zone of the larval olfactory pit, posterolaterally distributed, and more rounded in shape. Solitary ionocytes lacking *trpv6* and *ceacam1* were also seen in the non-sensory regions of the adult rosette. These observations suggest a complex mechanism for ion regulation throughout the zebrafish olfactory epithelium, involving region-specific roles for different ionocyte types.

**Figure 9.**
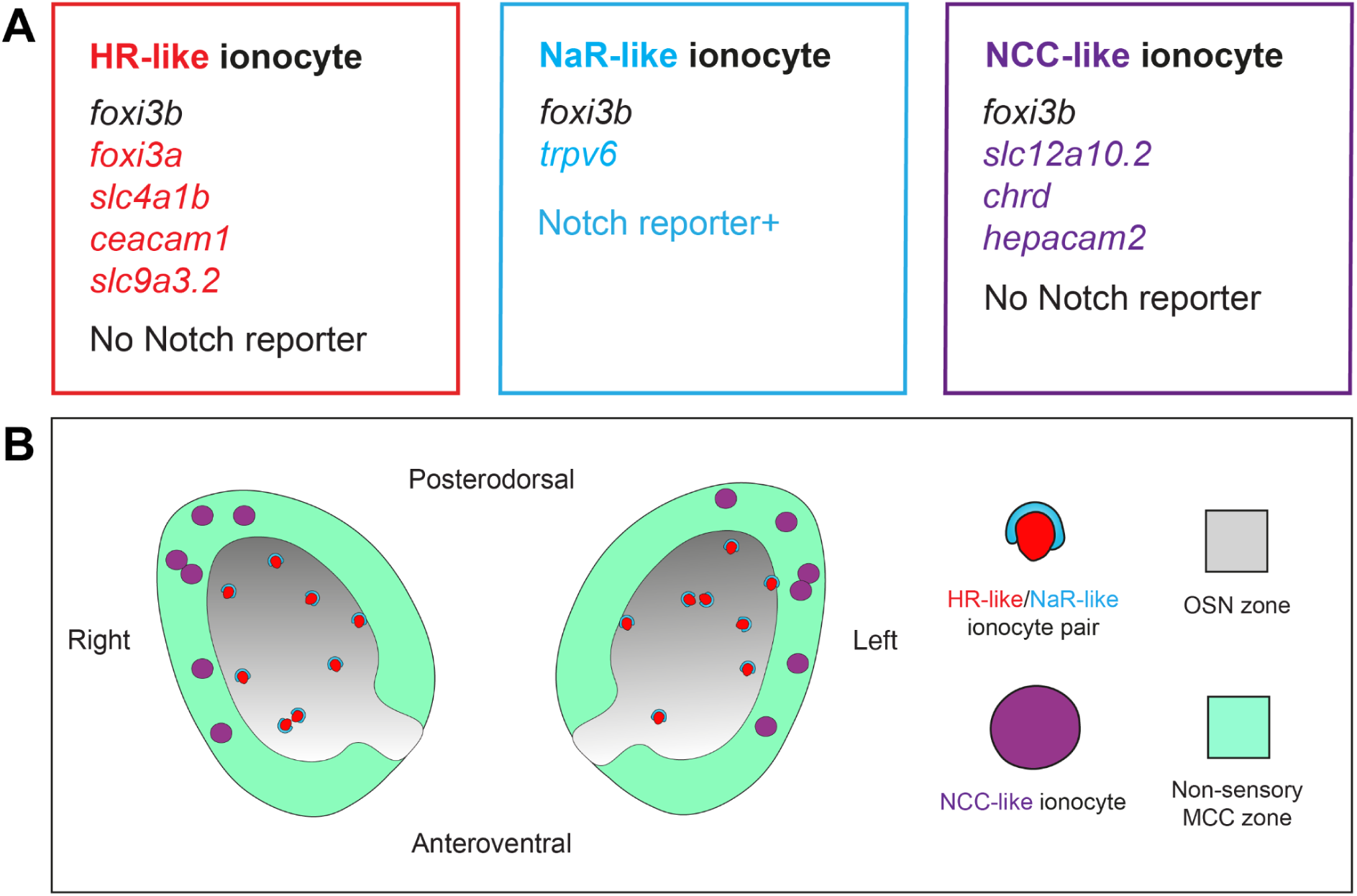
Summary of findings. **(A)** Gene expression in three different classes of olfactory ionocytes. **(B)** Schematic diagram showing the approximate number and distribution of ionocytes in the zebrafish larval olfactory pit (viewed from the front; not to scale).

**Figure 10.**
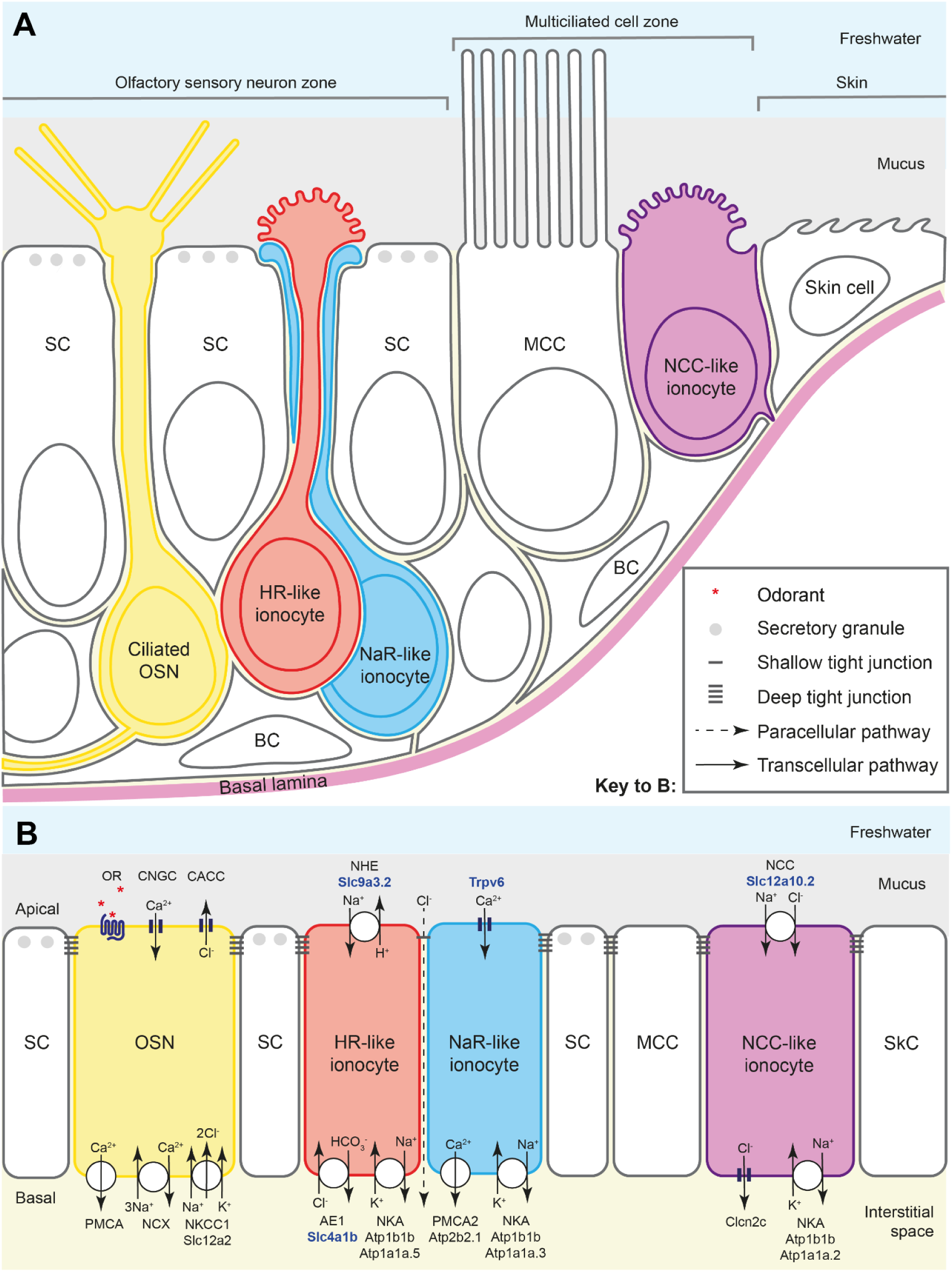
Ionocyte types and their proposed function in the zebrafish larval olfactory epithelium. **(A)** Schematic diagram to show the arrangement of HR- and NaR-like ionocyte pairs and NCC-like ionocytes in the zebrafish larval olfactory epithelium. The NCC-like cells have variable morphologies, some with a rounded microvillous apical surface. The diagram illustrates approximate cell shapes (not to scale). **(B)** Model for the contribution of olfactory ionocytes to ionic homeostasis in the zebrafish olfactory epithelium. This is a tentative proposal, based on mRNA expression data (see Figs. 1–4 and Supplementary Tables) and comparison to ionocytes in other tissues (see text for citations). The number and subcellular location of the relevant proteins in olfactory ionocytes have not yet been validated. Protein names in bold blue text are based on validated mRNA expression patterns in this study. Color coding as in panel A. The dotted line indicates the proposed paracellular pathway, via a shallow tight junction, between the HR-like and NaR-like olfactory ionocyte pair. This provides a route for recycling chloride ions, generated by sensory transduction in OSNs, within the OSN zone. **Abbreviations:** AE1 (Slc4a1b), anion exchanger 1; BC, basal cell; CACC, calcium-activated chloride channel; Clcn2c, chloride channel 2c; CNGC, cyclic-nucleotide-gated calcium channel; MCC, multiciliated cell; NCC (Slc12a10.2), sodium-chloride symporter; NCX, sodium-calcium exchanger; NHE (Slc9a3.2), sodium-proton exchanger; NKA (Atp1b1b, with cell-type-specific expression of alpha subunits), sodium-potassium ATPase; NKCC1 (Slc12a2), sodium-potassium-chloride cotransporter 1; OR, odorant receptor (GPCR), with odorant ligand (red asterisks); OSN, olfactory sensory neuron; PMCA, plasma membrane calcium ATPase; PMCA2 (Atp2b2.1), plasma membrane calcium ATPase 2; SC, supporting cell; SkC, skin cell; Trpv6, transient receptor potential vanilloid 6 (epithelial calcium channel).

### Pairing of NaR-like and HR-like ionocytes may extend functionality

NaR-like and HR-like ionocytes are consistently found as pairs in the olfactory epithelium. The cells are connected to each other by specialized tight junctions and are dependent on Notch signalling for survival, as can be seen by the loss of pre-existing paired ionocytes following treatment with LY411575 at 4 dpf. The existence of ionocytes in pairs or complexes is well established, for example in the gills and skin of stenohaline saltwater fish (Hootman and Philpott, 1980; Pisam and Rambourg, 1991), euryhaline fish adapted to seawater (Sardet et al., 1979; Karnaky, 1986; Shiraishi et al., 1997; Hiroi et al., 2005; Cozzi et al., 2015; Inokuchi et al., 2017), and in stenohaline freshwater teleosts (Hwang, 1988a). Pairs were also seen in the skin of the euryhaline medaka, when reared in freshwater (Hsu et al., 2014). In these cases, however, one member of the pair was termed an accessory cell.

Although accessory cells have some features of ionocytes — they are mitochondria-rich, for example — they have been considered to be immature or dormant (Hootman and Philpott, 1980) or specific to seawater fish (Sardet et al., 1979). The data here, and in the neuromast (Peloggia et al., 2021), indicate that both members of a pair in the freshwater zebrafish are *bona fide* ionocytes.

The conservation of architecture across evolutionarily diverse fish species suggests a functional advantage in using pairs. One consequence of ionocyte pairing is the creation of a paracellular pathway, which enables the movement of ions down an electrochemical gradient. The permeability of the paracellular pathway is dependent on Claudins, with anion transport enabled by Claudin-4 in tandem with Claudin-8 (Hou et al., 2010), as well as Claudin-17 (Krug et al., 2012). In the olfactory epithelium, the apical half of the NaR-like cell wraps around the HR-like cell (Fig. 10A), forming a paracellular pathway in the space between the two cells (Fig. 10B). Both members of the pair extend from just above the basal lamina to the surface within the sensory zone. Here, it is possible that the ionocyte pairs directly regulate ion concentration within the mucus and epithelium to optimize olfactory transduction. For example, one possible role could be to enable uptake of Cl^-^ after a neuronal signalling event, with apical uptake and basal secretion of Na^+^ by NaR-like and HR-like ionocytes providing the driving force (Fig. 10B). Consistent with anion transport via a paracellular pathway, olfactory ionocytes appear to express *cldn17* (https://daniocell.nichd.nih.gov/gene/C/cldn17/cldn17_tast.html∼). Intriguingly,

NaR-like/HR-like ionocyte pairs are also found in the zebrafish neuromast (Peloggia et al., 2021), which contains sensory hair cells that – like OSNs – signal with a chloride efflux (Lunsford et al., 2023). Thus, this specific pair may enable transport of ions that are required for this mode of signalling.

### NCC-like olfactory ionocytes are located in non-sensory regions containing motile cilia

NCC-like ionocytes are strikingly different from NaR-like and HR-like ionocytes in a number of respects. In the larval and adult olfactory epithelium, NCC-like ionocytes appear to be restricted to non-sensory regions. Here, they are interspersed among multiciliated cells, which contain motile cilia that drive a constant water flow over the olfactory pit (Cox, 2008; Reiten et al., 2017). By contrast, NCC ionocytes are absent from the zebrafish neuromast, which lack motile cilia. A close association between ionocytes and motile cilia has been observed in the skin of *Xenopus tropicalis*, where depletion of ionocytes disrupts ciliary beating (Dubaissi and Papalopulu, 2011). In the mammalian airway, ciliary beat is regulated by intracellular chloride levels (Inui et al., 2019; Yasuda et al., 2020). NCC-like ionocytes can potentially transport chloride via Slc12a10.2 and Clcn2c (Pérez-Rius et al., 2015) (Fig. 10B; see also Table S2). These observations raise the possibility that NCC-like ionocytes function specifically to influence ciliary beating in the zebrafish olfactory epithelium, which in turn shapes the detection of odorants (Reiten et al., 2017).

### Shared and unique properties of zebrafish olfactory ionocytes

In mammals, Foxi1^+^ ionocytes have been identified in various tissues including the kidney (Rao et al., 2019), airway epithelium (Montoro et al., 2018; Yuan et al., 2023), inner ear (Honda et al., 2017), salivary gland (Mauduit et al., 2022) and thymus (Bautista et al., 2021). These ionocytes, which are critical to organ function, display a number of tissue-specific properties, including type of transporters expressed and morphology. Tissue-specific gene expression in zebrafish ionocytes is illustrated by the larval transcriptome: olfactory ionocytes cluster with olfactory epithelial cells, rather than skin ionocytes. The clustering of ionocytes within their tissue of residency indicates that they share a transcriptional signature with the cells they regulate. Some of these genes are likely to code for adhesion molecules, which are highly enriched in all ionocyte transcriptomes that we have analysed thus far. Other genes may be involved in cell-cell communication and cell signalling pathways.

In summary, this study has yielded two unexpected features of ionocytes in the zebrafish olfactory epithelium. Firstly, the epithelium contains pairs of NaR-like/HR-like ionocytes, implying that synergy between these ionocyte types is essential for maintaining ion balance. Secondly, NCC-like ionocytes are restricted to non-sensory regions, implying a distinct function in the regulation of motile cilia. Ionocytes thus have diverse roles in enabling optimal olfaction.

## Acknowledgements

We are grateful to Cynthia Shiyuan Chen for her help with scRNA-seq data analysis and to Nathalie Jurisch-Yaksi for generous discussion of the adult olfactory transcriptome datasets at early stages of this study. We thank Chris Hill for technical assistance with Scanning Electron Microscopy, which was carried out in the Cryo-Electron Microscopy Facility, University of Sheffield.

## Author Contributions

Conceptualization: JP, TP, TTW, SJ; Data Analysis: JP, KYC, TTW; Funding acquisition: FE, JWL, TP, TTW, SJ; Investigation: JP, KYC, TTW, MDP, RS, JBW, YW, SW, MJ, SJ; Methodology: YW, SW, MJ, JWL, FE, VJ ; Resources: TP, TTW, JWL, FE, VJ, MJ, SJ; Supervision: JWL, FE, VJ, SJ, TTW, TP; Validation: JP; Visualization: JP, NvH, TTW; Writing — original draft: JP, KYC, TTW, SJ; Writing — review and editing: SJ and TTW, with input from MDP, TP and KYC.

## Data availability

Raw data underlying this manuscript can be accessed from the Stowers Original Data Repository at http://www.stowers.org/research/publications/libpb-2509.

## Licencing

For the purpose of open access, the authors will apply a Creative Commons Attribution Licence (CC BY) to any Author Accepted Manuscript arising from this study.

---

**Figure S1, related to Figure 1.**
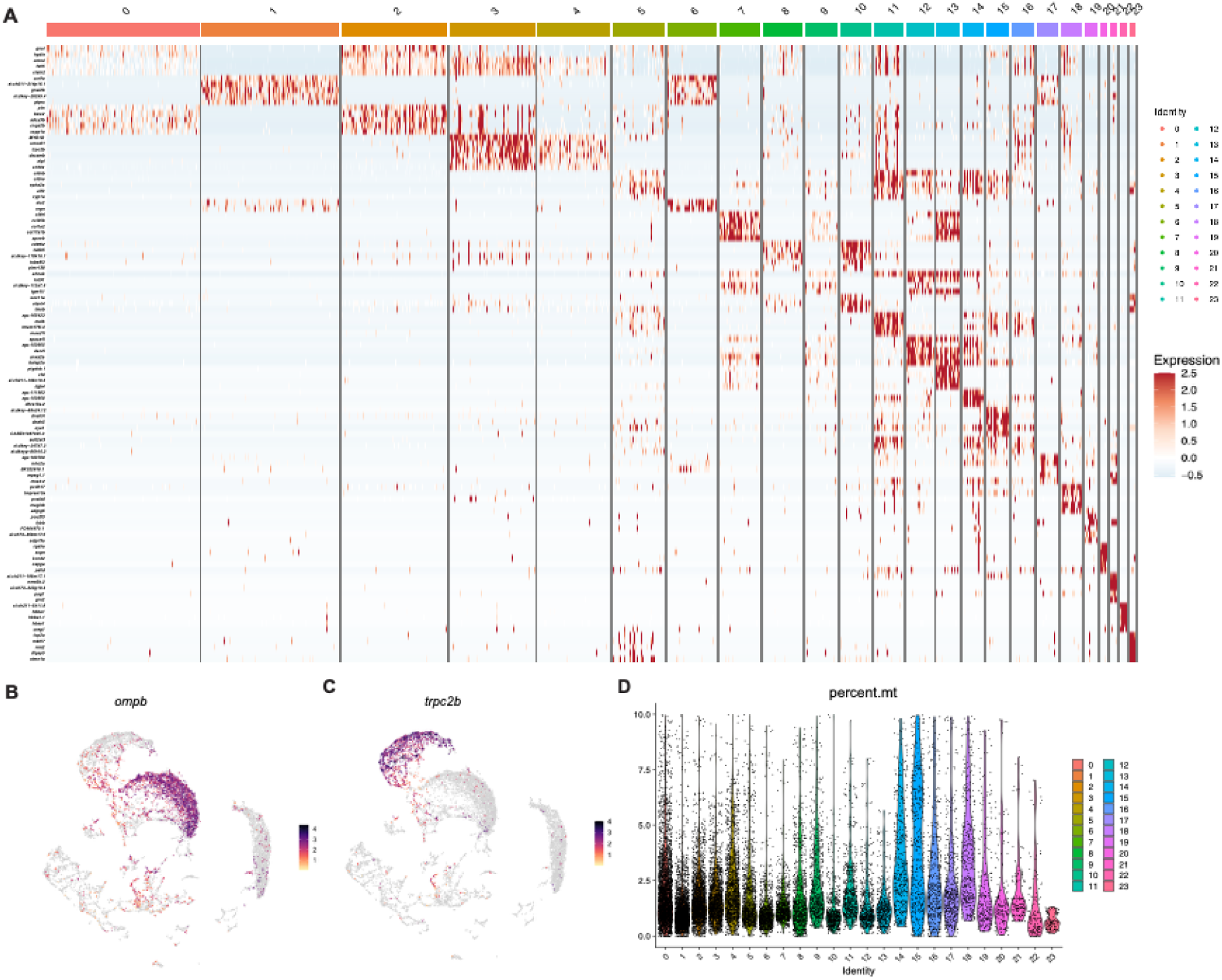
**(A)** Heatmap showing top 5 highest expressing genes based on LogFC on the clusters obtained in the dataset from dissected adult zebrafish olfactory organs. Based on these genes and markers given previously (Kraus et al., 2022), the clusters can be classified as: ciliated neurons: clusters 0, 2; microvillous neurons: clusters 3, 4; neuronal precursors: clusters 8, 10, 23; early progenitors: clusters 14, 15; sustentacular cells: clusters 7, 9, 12, 13, 20; immune cells: clusters 1, 6, 11, 17, 21. **(B)** Feature plots of *trpc2b* (microvillous OSN marker) and **(C)** *ompb* (ciliated OSN marker). **(D)** Violin plot showing the percentage of mitochondrial genes in the dataset.

**Figure S2, related to Figure 3.**
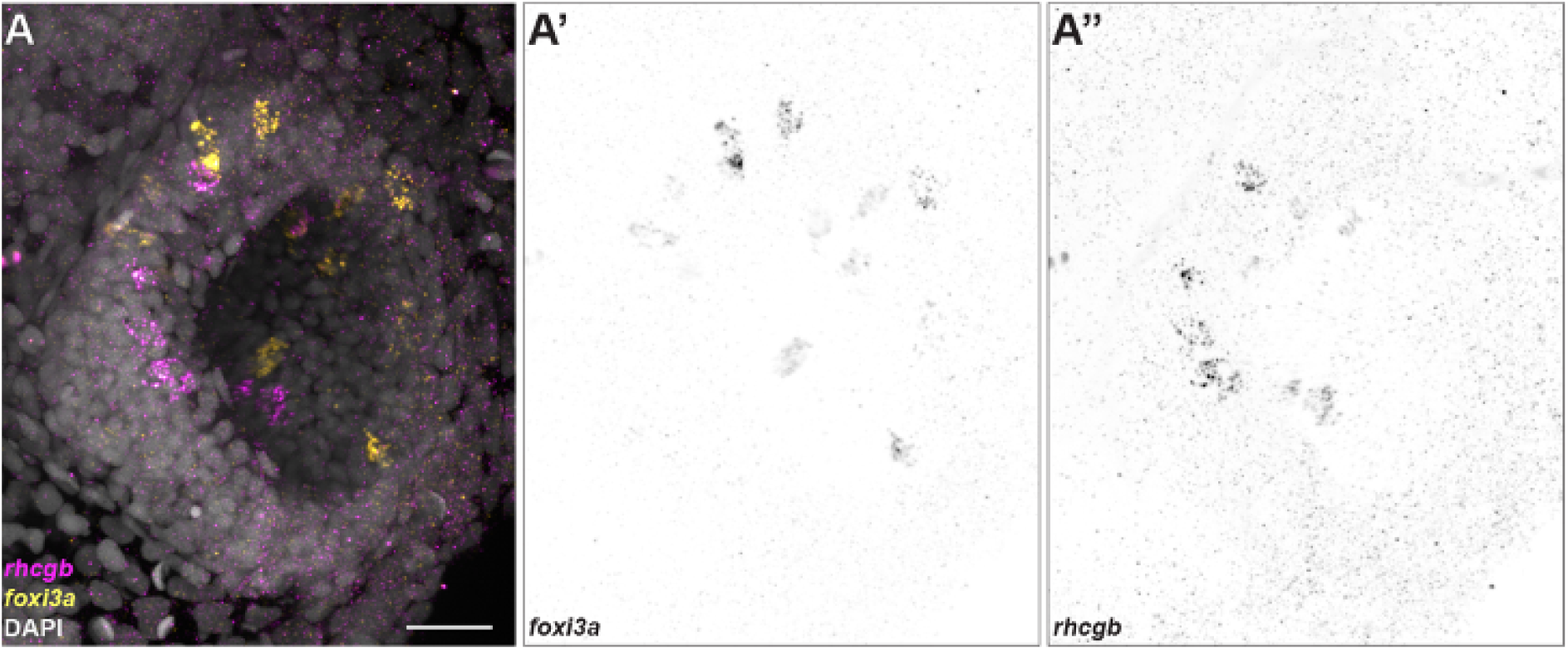
**(A)** Maximum intensity projection of a confocal image showing HCR RNA-FISH signals for *foxi3a* (yellow) and *rhcgb* (magenta) with DAPI stain (grey). **(A’)** Individual channels for *foxi3a* and **(A’’)** *rhcgb*. Scale bar: 20 µm.

**Figure S3, related to Figure 4.**
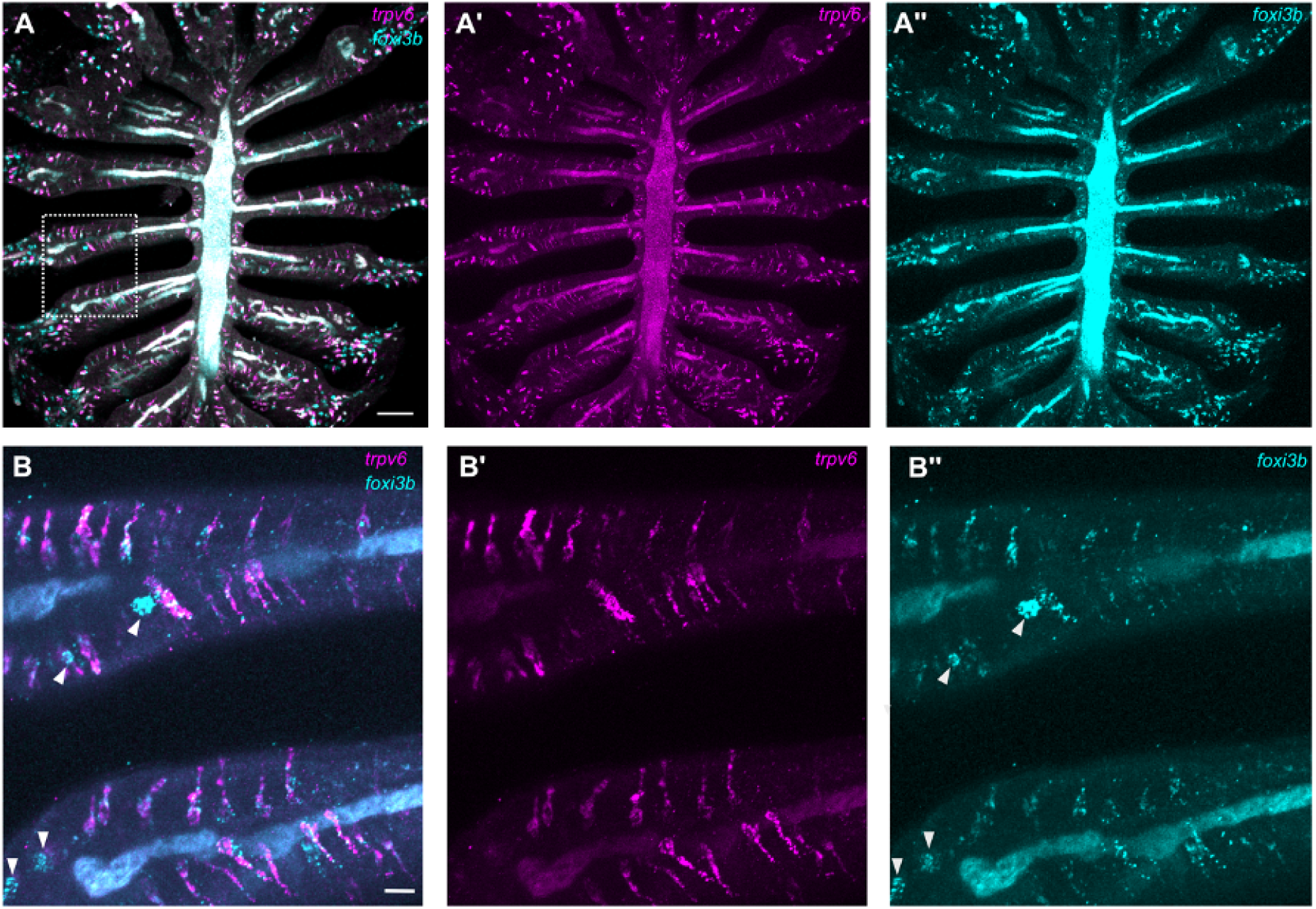
**(A)** Overview of an adult olfactory rosette, showing expression of *trpv6* **(A’)** and *foxi3b* **(A’’)**. **(B–B’’)** High magnification view of the region outlined in panel A. The arrowheads indicate solitary NCC-like ionocytes, which express *foxi3b* and have a rounded shape. These are distinct from elongated pairs of cells that express either *foxi3b* (cyan) or *trpv6* (magenta). Scale bar: **A**, 50 µm; **B**, 20 µm.

**Figure S4, related to Figure 5.**
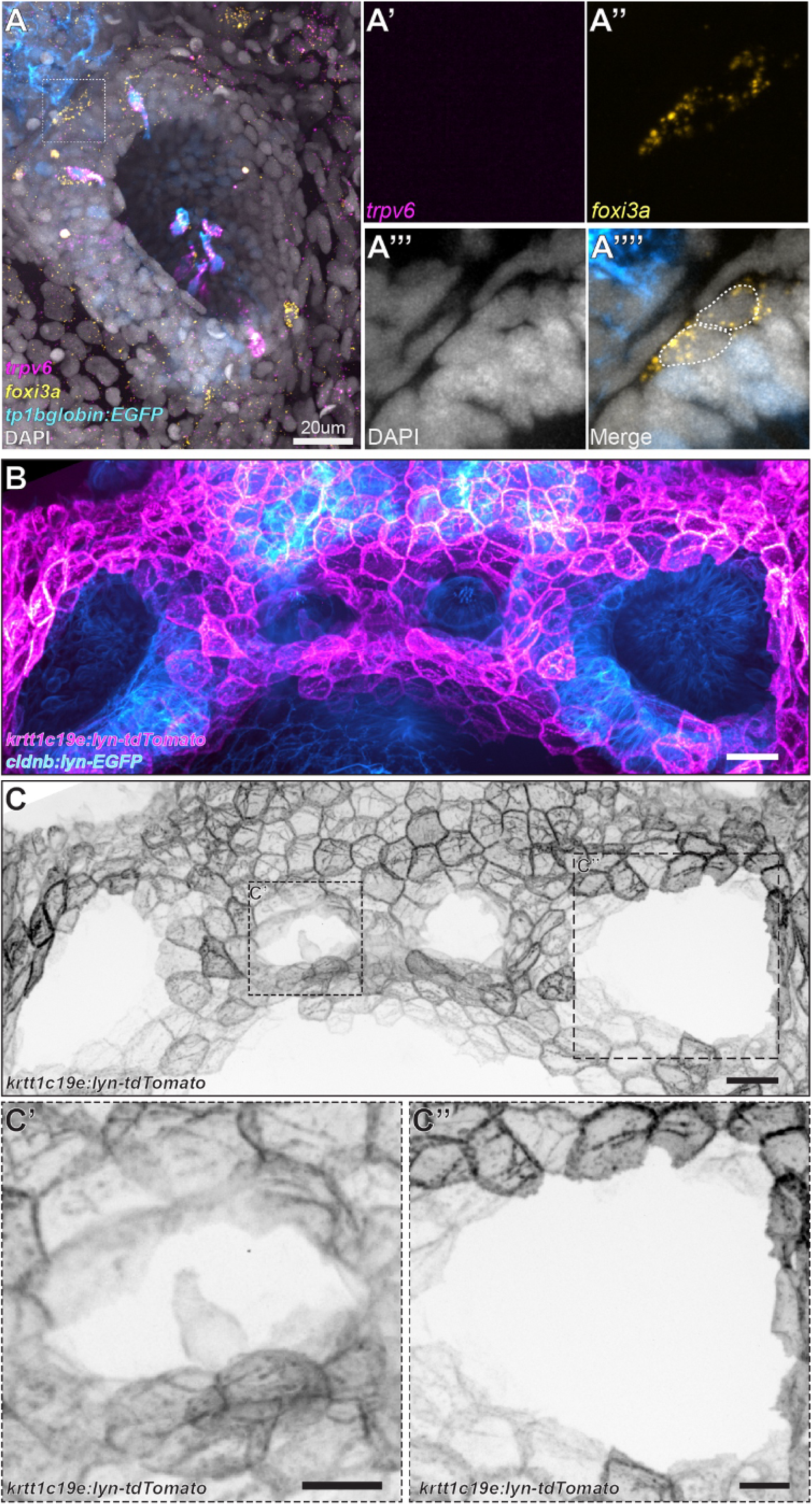
**(A–A’’’’)** *foxi3a^+^* cells (yellow) at the edge of an olfactory pit, as shown by HCR RNA-FISH. **(B)** Maximum intensity projection of a confocal image from *Tg(-8.0cldnb:lyn-EGFP)^zf106Tg^;Tg(krtt1c19e:lyn-tdTomato)^sq16^*transgenic larva shows a pair of Nm ionocytes in the neuromast, but no tdTomato^+^ cells in the olfactory pit. **(C)** Single channel image of *krtt1c19e:lyn-tdTomato*. **(C’)** Enlargement of a neuromast containing tdTomato^+^ Nm ionocytes. **(C’’)** Enlargement of an olfactory pit containing no tdTomato^+^ cells. Scale bars: **A**, 20 µ; **B–C’’**, 5 µm.

## Supplementary Material

**Supplementary Table 1. Differentially expressed genes in dissected adult zebrafish olfactory organ scRNA-seq dataset**

This spreadsheet contains the differentially expressed gene list (cluster markers) of olfactory cell types from the integrated adult dataset via Seurat::FindAllMarkers function with default parameters. Ionocytes are on cluster 18.

**Supplementary Table 2. Differentially expressed genes in larval 5 dpf olfactory cell subset**

This spreadsheet contains the differentially expressed gene list (cluster markers) of olfactory cell types from the subsetted larval dataset via Seurat::FindAllMarkers function with default parameters.

**Movie 1. 3D rendering of an embryonic olfactory pit depicting HR- and NaR-like ionocytes**

HCR RNA-FISH for *trpv6* (magenta) and *foxi3a* (yellow), combined with the Notch reporter *tp1bglobin:EGFP* (cyan), shows spatial distribution of ionocyte pairs in the larval 5 dpf olfactory pit. Initial image is a frontal view of the left olfactory pit, with dorsal to the top and lateral to the right. Scale bar: 15 µm.

**Movie 2. 3D rendering of an embryonic olfactory pit showing NCC-like ionocytes**

HCR RNA-FISH for *slc12a10.2* (yellow) and *foxi3b* (magenta), combined with the Notch reporter *tp1bglobin:EGFP* (cyan), showing the spatial distribution of ionocyte pairs in the larval 5 dpf olfactory pit. Initial image is a frontal view of the left olfactory pit, with dorsal to the top and lateral to the right. Scale bar: 15 µm.

**Movie 3. New olfactory ionocytes do not express skin transgenes**

Time lapse from *Tg(-8.0cldnb:lyn-EGFP)^zf106Tg^;Tg(krtt1c19e:lyn-tdTomato)^sq16^*transgenic zebrafish larva showing pairs of tdTomato^+^ Nm ionocytes (arrow, middle panel) invading neuromasts, but no positive cells in the olfactory epithelium. Scale bar: 5 µm.

**Movie 4. Differentiation of ionocyte pairs in a larval olfactory pit and neuromast**

Time lapse from a 3 dpf transgenic zebrafish larva (*Tg(dld:hist2h2l-EGFP)^psi84^*) showing a pair of ionocytes (cyan and yellow dots; visible at the start of the recording) invading the neuromast. At ∼90 hours ionocytes are visible in the olfactory pit (red and cyan dots). No invasion was detected. Scale bar: 5 µm.

**Movie 5. Ionocyte pair development in a larval olfactory pit**

Time lapse from a 3 dpf transgenic zebrafish larva (*Tg(dld:hist2h2l-EGFP)^psi84^*) showing the appearance of a pair of olfactory ionocytes (red and cyan dots). The pair is visible at approximately 85 hours, and move around together. Scale bar: 5 µm.

**Movie 6. 3D reconstruction of an HR-like/NaR-like ionocyte pair from a 7 dpf wild-type olfactory pit**

360° rotation of Fig. 7A. Red, HR-like cell; cyan, NaR-like cell.

**Movie 7. 3D reconstruction of the tight junctions of an HR-like/NaR-like ionocyte pair**

360° rotation of Fig. 7G; compare to Movie 9. Green, shallow tight junction between the two ionocytes; yellow, deep tight junction between NaR-like ionocyte and olfactory supporting cell; orange, deep tight junction between NaR-like ionocyte and multiciliated cell; blue, deep tight junction between HR-like ionocyte and olfactory supporting cell.

**Movie 8. 3D reconstruction of a 3-cell ionocyte complex from a 7 dpf wild-type olfactory pit**

360° rotation of Fig. 7L. Red, HR-like cell; cyan, NaR-like cell; dark blue, possible second NaR-like cell; yellow, ciliated OSN (not part of the ionocyte complex, but included for context and scale).

**Movie 9. 3D reconstruction of the tight junctions between the HR-like (red) and NaR-like (cyan) ionocytes in Fig. 7L**

360° rotation of Fig. 7P; compare to Movie 7. Green, shallow tight junction between the two ionocytes; yellow, deep tight junction between NaR-like ionocyte (cyan) and olfactory supporting cell; blue, deep tight junction between HR-like ionocyte (red) and olfactory supporting cell.

