## Supplementary Figures for "Paired and solitary ionocytes in the zebrafish olfactory epithelium"

### Supplementary Material

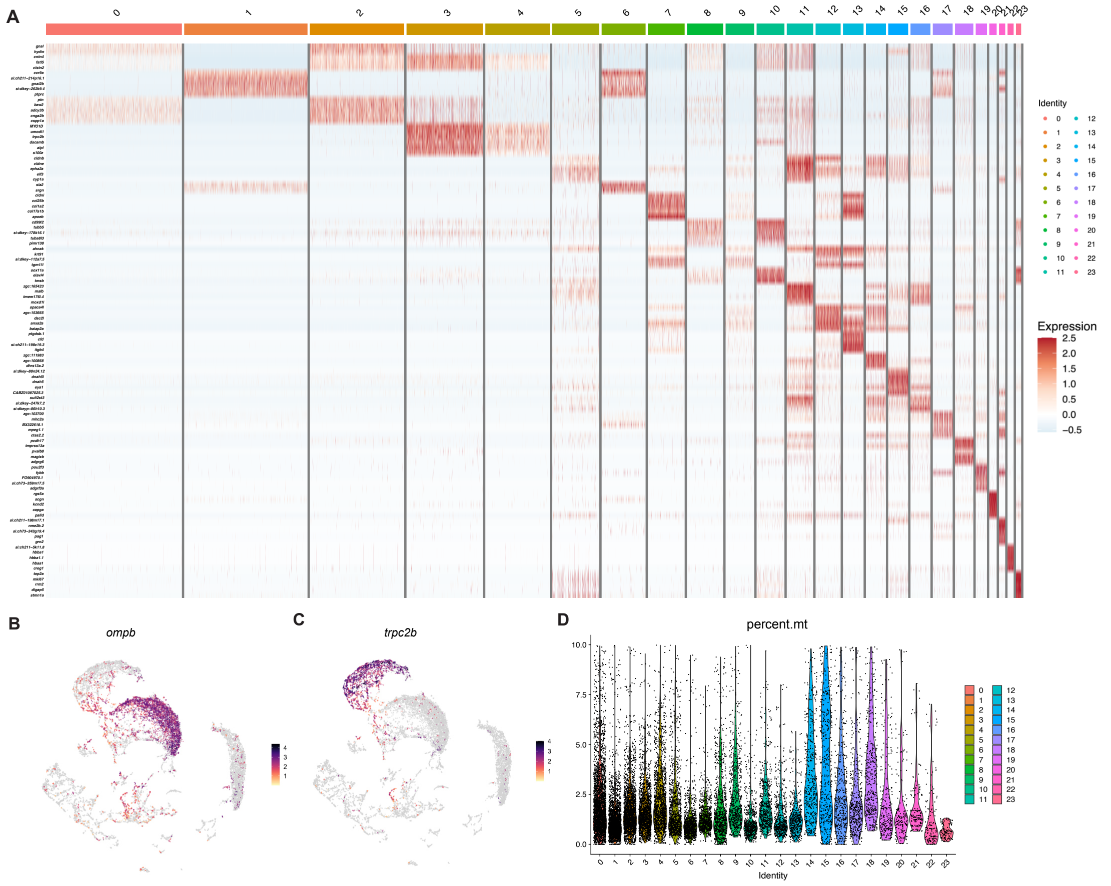


**Figure S1, related to Figure 1**

**(A)** Heatmap showing top 5 highest expressing genes based on LogFC on the clusters obtained in the dataset from dissected adult zebrafish olfactory organs. Based on these genes and markers given previously (Kraus et al., 2022), the clusters can be classified as: ciliated neurons: clusters 0, 2; microvillous neurons: clusters 3, 4; neuronal precursors: clusters 8, 10, 23; early progenitors: clusters 14, 15; sustentacular cells: clusters 7, 9, 12, 13, 20; immune cells: clusters 1, 6, 11, 17, 21. **(B)** Feature plots of *trpc2b* (microvillous OSN marker) and **(C)** *ompb* (ciliated

OSN marker). **(D)** Violin plot showing the percentage of mitochondrial genes in the dataset.

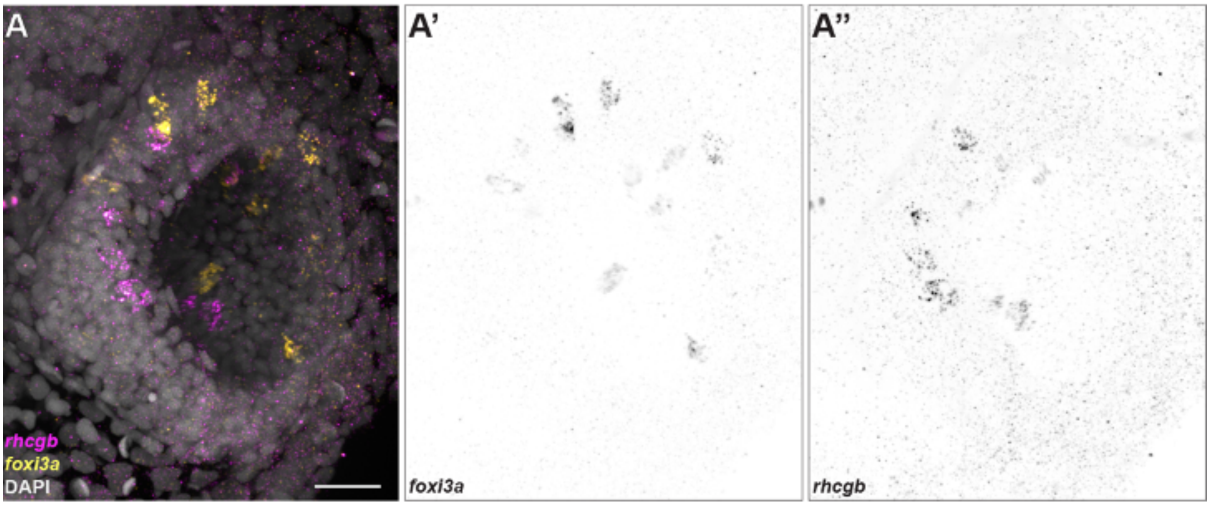


**Figure S2, related to Figure 3**

**(A)** Maximum intensity projection of a confocal image showing HCR RNA-FISH signals for *foxi3a* (yellow) and *rhcgb* (magenta) with DAPI stain (grey). **(A')** Individual channels for *foxi3a* and **(A'')** *rhcgb*. Scale bar: 20 μm.

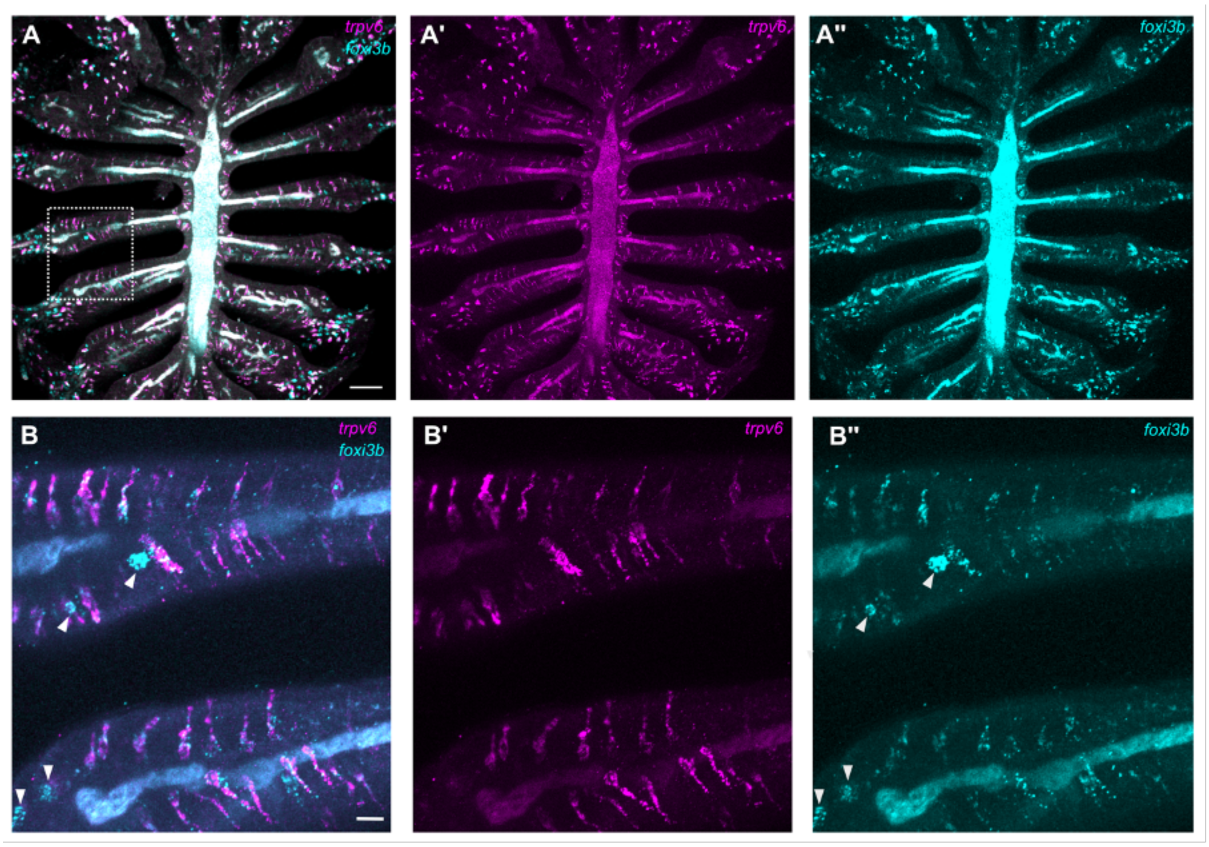


**Figure S3, related to Figure 4**

**(A)** Overview of an adult olfactory rosette, showing expression of *trpv6* **(A')** and *foxi3b* **(A'')**. **(B–B'')** High magnification view of the region outlined in panel A. The arrowheads indicate solitary NCC-like ionocytes, which express *foxi3b* and have a rounded shape. These are distinct from elongated pairs of cells that express either *foxi3b* (cyan) or *trpv6* (magenta). Scale bar: **A**, 50 µm; **B**, 20 µm.

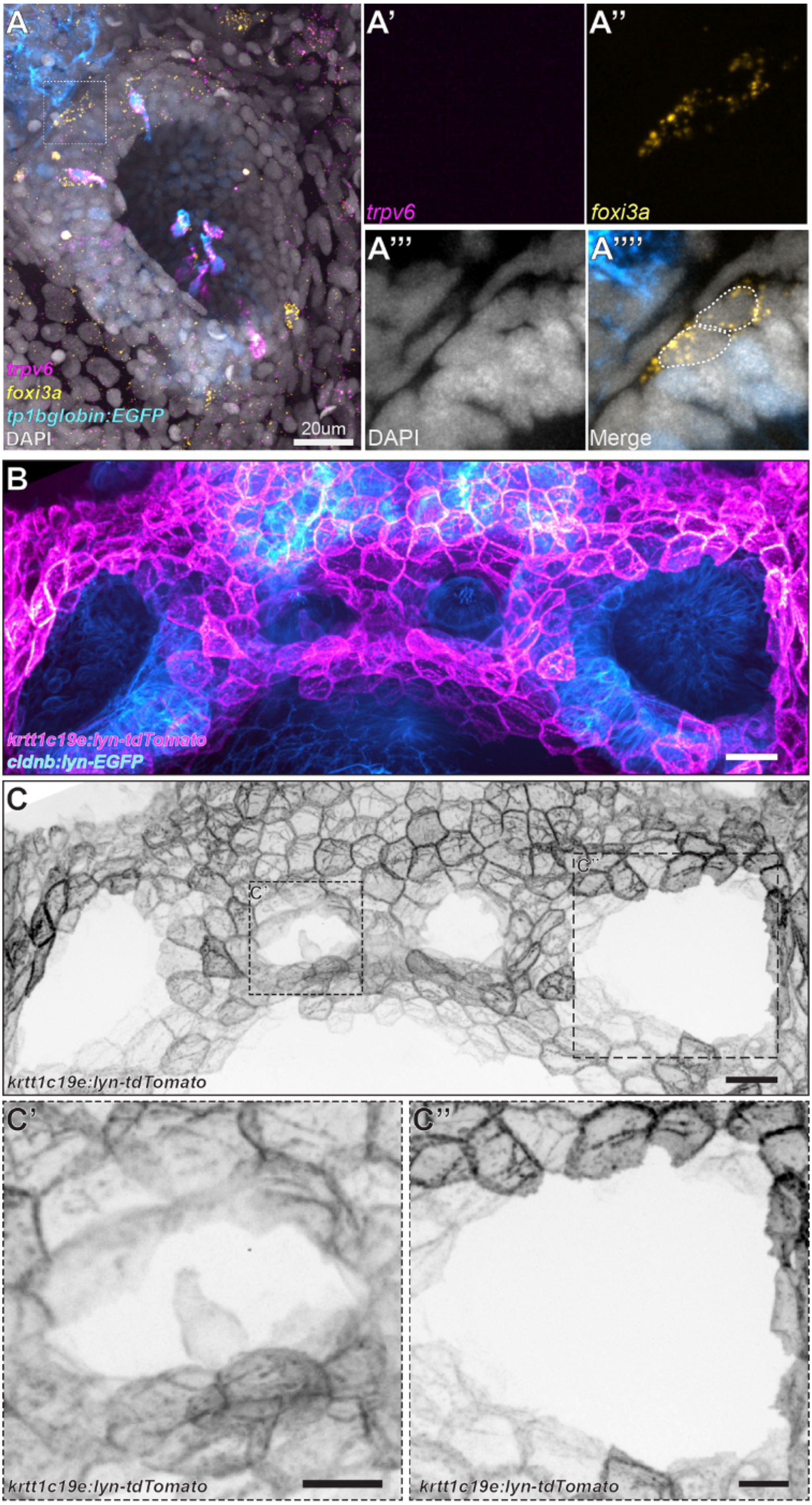
A
A'
A''
trpv6
foxi3a
A'''
A''''
trpv6
foxi3a
tp1bglobin:EGFP
DAPI
20um
DAPI
Merge
B
krtt1c19e:lyn-tdTomato
cldnb:lyn-EGFP
C
C'
C''
krtt1c19e:lyn-tdTomato
C'
C''
krtt1c19e:lyn-tdTomato
krtt1c19e:lyn-tdTomato

**Figure S4, related to Figure 5**

**(A–A'''')** *foxi3a*$^{+}$ cells (yellow) at the edge of an olfactory pit, as shown by HCR RNA-FISH. **(B)** Maximum intensity projection of a confocal image from *Tg(-8.0cldnb:lyn-EGFP)*$^{zf106Tg}$*;Tg(krtt1c19e:lyn-tdTomato)*$^{sq16}$ transgenic larva shows a pair of Nm ionocytes in the neuromast, but no tdTomato$^{+}$ cells in the olfactory pit. **(C)** Single channel image of *krtt1c19e:lyn-tdTomato*. **(C')** Enlargement of a neuromast containing tdTomato$^{+}$ Nm ionocytes. **(C'')** Enlargement of an olfactory pit containing no tdTomato$^{+}$ cells. Scale bars: **A**, 20 μ; **B–C''**, 5 μm.
